# Olfactory facilitation of visual categorization in the 4-month-old brain depends on visual demand

**DOI:** 10.1101/2024.06.09.598135

**Authors:** Anna Kiseleva, Diane Rekow, Benoist Schaal, Arnaud Leleu

## Abstract

To navigate their environment, infants rely on intersensory facilitation when unisensory perceptual demand is high, a principle known as inverse effectiveness. Given that this principle was mainly documented in the context of audiovisual stimulations, here we aim to determine whether it applies to olfactory-to-visual facilitation. We build on previous evidence that the mother’s body odor facilitates face categorization in the 4-month-old brain, and investigate whether this effect depends on visual demand. Scalp electroencephalogram (EEG) was recorded in 2 groups of 4-month-old infants while they watched 6-Hz streams of visual stimuli with faces displayed every 6^th^ stimulus to tag a face-selective response at 1 Hz. We used variable natural stimuli in one group (*Nat* Group), while stimuli were simplified in the other group (*Simp* Group) to reduce perceptual categorization demand. During visual stimulation, infants were alternatively exposed to their mother’s *vs*. a baseline odor. For both groups, we found an occipito-temporal face-selective response, but with a larger amplitude for the simplified stimuli, reflecting less demanding visual categorization. Importantly, the mother’s body odor enhances the response to natural, but not to simplified, face stimuli, indicating that maternal odor improves face categorization when it is most demanding for the 4-month-old brain. Overall, this study demonstrates that the inverse effectiveness of intersensory facilitation applies to the sense of smell during early perceptual development.

**Highlights:**

- Intersensory facilitation is a function of unisensory perceptual demand in infants (inverse effectiveness).
- This inverse relation between multisensory and unisensory perception has been mainly documented using audiovisual stimulations.
- Here we show that olfactory-to-visual facilitation depends on visual demand in 4-month-old infants.
- The inverse effectiveness of intersensory facilitation during early perceptual development applies to the sense of smell.

## 1. Introduction

Human infants are exposed to a complex multisensory environment from the very beginning of life. The early ability to form multisensory percepts has long been debated (e.g., Gibson, 1969; Piaget, 1952), but evidence that infants’ sensory systems interact is nowadays clear (Bahrick and Lickliter, 2012; Lewkowicz and Bremner, 2020; Murray et al., 2016; Streri and de Hevia, 2023, for reviews). Newborns already possess the brain architecture to integrate inputs across sensory modalities (Sours et al., 2017) and have incipient skills to respond to correlated multisensory cues, relying on intensity (Lewkowicz and Turkewitz, 1980) or temporal occurrence (Lewkowicz et al., 2010). Multisensory perception then develops during the first months of life, allowing infants to bind sensory inputs insofar as they co-occur in space (Neil et al., 2006) and/or time (e.g., Hyde et al., 2011; Lewkowicz, 1996; Werchan et al., 2018).

An influential view of multisensory development, the intersensory redundancy hypothesis, assumes that infants rely on the detection of redundant amodal properties (e.g., rhythm, intensity, synchrony) across sensory inputs to integrate them (Bahrick and Lickliter, 2000). Interestingly, this view predicts that, at early developmental stages, multisensory stimuli are perceived more efficiently than unisensory stimuli, namely *intersensory facilitation* (Bahrick and Lickliter, 2012). As a result, infants can discriminate the tempo of synchronized auditory and visual inputs at 3 months (Bahrick et al., 2002), their rhythm at 5 months (Bahrick and Lickliter, 2000), or their prosody (Bahrick et al., 2019) and emotion (Flom and Bahrick, 2007) at 4 months, whereas they cannot with unisensory or asynchronous inputs. Later on, as sensory systems develop, unisensory and multisensory stimulations gradually become equally effective, whereas intersensory facilitation declines. For example, while 3- and 5-month-old infants discriminate the tempo and rhythm of an audiovisual stimulation, respectively, they fail when exposed to the sole visual input (Bahrick et al., 2002; Bahrick and Lickliter, 2000). In contrast, at 5 and 8 months of age for tempo and rhythm, respectively, infants discriminate both unisensory (visual) and multisensory (audiovisual) conditions, with no more intersensory facilitation (Bahrick and Lickliter, 2004).

These results suggest that intersensory facilitation in early development follows a well-known principle of multisensory integration called *inverse effectiveness* (e.g., Meredith and Stein, 1983; Stevenson et al., 2007). According to this principle, the weakest unisensory responses lead to the strongest multisensory integration. In human adults, inverse effectiveness was established by increasing task load (e.g., Santangelo and Spence, 2007) or decreasing the effectiveness of unisensory stimuli to trigger a response, whether it is behavioral (Regenbogen et al., 2016), hemodynamic (Stevenson et al., 2007; Stevenson and James, 2009), or electroencephalographic (EEG) (Stevenson et al., 2012). Similarly, in infants, the strength of intersensory facilitation relates to the effectiveness of unisensory discrimination (Bahrick et al., 2010). The authors tested 5-month-old infants and used a tempo discrimination task that is usually achieved at that age with both visual and audiovisual stimulations and without any intersensory facilitation (Bahrick & Lickliter, 2004). By increasing the difficulty of the task, they found that intersensory facilitation reappears while unisensory discrimination disappears. This indicates that intersensory facilitation not only declines as unisensory perception develops, but also depends on unisensory *perceptual demand* at a given age.

With the notable exception of touch (e.g., Ronga et al., 2021), inverse effectiveness in infancy was mostly investigated in the auditory and visual domains. Thus, whether this principle generalizes to other sensory realms remains an open question. In particular, the sense of smell has some specificities compared to the other senses that directly question whether it relies on the same perceptual mechanisms. A main difference is the sensitivity to spatiotemporal synchrony, which is poor for olfaction as opposed to vision and audition because the olfactory system goes beyond the spatial variability introduced by the volatility of odorant molecules and the temporal discontinuity imposed by breathing to perceive odors as more constant than they are in reality (Sela and Sobel, 2010). However, despite this loose spatiotemporal sensitivity, earlier studies have shown intersensory facilitation involving olfaction in infants (e.g., Durand et al., 2013; Fernandez and Bahrick, 1994; Godard et al., 2016; Jessen, 2020; reviewed in Damon et al., 2021; Schaal and Durand, 2012). For instance, recent EEG studies in 4-month-old infants found that the rapid categorization of naturalistic face stimuli against a variety of nonface stimuli, which represents a challenging visual task for the infant brain, is facilitated when the mother’s body odor is presented concurrently (Leleu et al., 2020; Rekow et al., 2020). At the same age, maternal odor even initiates face-selective neural activity in response to ambiguous face-like objects (Rekow et al., 2021). Interestingly, in line with aforementioned audiovisual studies, intersensory facilitation by the mother’s odor gradually declines as rapid face categorization develops between 4 and 12 months (Rekow et al., 2023). Later on, in adults exposed to both genuine human faces and ambiguous face-like objects, only the EEG response to face-like objects is enhanced by a concomitant body odor, human faces being readily categorized through vision alone whereas face-like objects are less effectively categorized (Rekow et al., 2022). This developmental trade-off suggests that olfactory-to-visual facilitation, like audiovisual facilitation, follows the inverse effectiveness principle as it progressively disappears when visual categorization improves and becomes effective on its own.

Here, to directly evaluate whether the inverse effectiveness principle applies to olfactory-to-visual facilitation at a young age, we built upon previous audiovisual studies that manipulated perceptual demand (e.g., Bahrick et al., 2010) and used a frequency-tagging EEG approach to measure the maternal odor effect on rapid face categorization in two groups of 4-month-old infants. The first group (*Nat* Group) was exposed to a highly variable set of *natural* face and nonface stimuli as used in previous studies showing the effect (Leleu et al., 2020; Rekow et al., 2023, 2021, 2020). The second group (*Simp* Group) was exposed to a less variable set of *simplified* stimuli to reduce physical variability across face stimuli while keeping high variability between face and nonface stimuli, thus decreasing perceptual demand compared to natural stimuli. In both groups, stimuli were presented as fast streams of 6 stimuli/s (at 6 Hz) with faces inserted every 6^th^ stimulus to tag a face-selective response at 1 Hz and harmonics (i.e., integer multiples) in the EEG spectrum. During visual stimulation, infants were alternatively exposed to baseline or a maternal odor context. We expected a larger face-selective response in Simp Group than in Nat Group, reflecting less demanding face categorization for the former. We also expected a larger EEG response in the maternal than in the baseline odor context only for Nat Group, indicating olfactory-to-visual facilitation when visual perception is not fully effective.

## 2. Material and methods

### 2.1. Participants

Forty-eight full-term White 4-month-old infants recruited through the local birth registry, and reported by parents as being in optimal general heath without any sensory (e.g., visual, olfactory) or neurological disorder, participated in the study. Before testing, parents were informed about the objectives and methods of the study and signed a written informed consent. Testing was conducted according to the Declaration of Helsinki and approved by the French ethics committee (CPP Sud-Est III - 2016-A02056-45). Data from six infants were excluded on the basis of less than two valid sequences per odor context (*N* = 3), or too noisy EEG data (*N* = 3). The final sample was thus composed of 42 infants who were randomly grouped according to the type of visual stimuli they were exposed to (see section 2.2): 21 infants were exposed to natural stimuli (Nat Group: 9 females, mean age ± *SD*: 135 ± 13 days, range: 112–168 days) and 21 infants to simplified stimuli (Simp Group: 9 females, mean age ± *SD*: 132 ± 10 days, range: 119–162 days). The two groups did not significantly differ in age (*T* (40) = 0.91, *p* = .37). Sample size was estimated a priori from the maternal odor effect found in Leleu et al. (2020) that we aimed to replicate in Nat Group in the present study. We considered a Cohen’s *d* = +0.79 (effect size at right occipito-temporal electrodes), a significance level *α* = .05 (two-tailed), and a high power 1-*β* = .95 to optimize our ability to replicate the effect, leading to a sample size *N* = 21. For the sake of simplicity, we equalized sample size in both groups.

### 2.2. Visual stimuli

Visual stimuli consisted in pictures of human adult faces and various living and non-living objects (i.e., animals, plants, man-made objects) cropped to a square and resized to 400 × 400 pixels. Two stimulus sets were created (one for each group of infants), each composed of 68 faces (34 females) and 170 objects (85 categories × 2 exemplars; see examples in Figure 1A; full stimulus sets in Figure S1). Stimuli were adapted from previous studies (e.g., Leleu et al., 2020; Liu-Shuang et al., 2014; Rekow et al., 2023), from the Face Research Lab London Set (DeBruine and Jones, 2017) or collected from the Internet.

**Figure 1.**
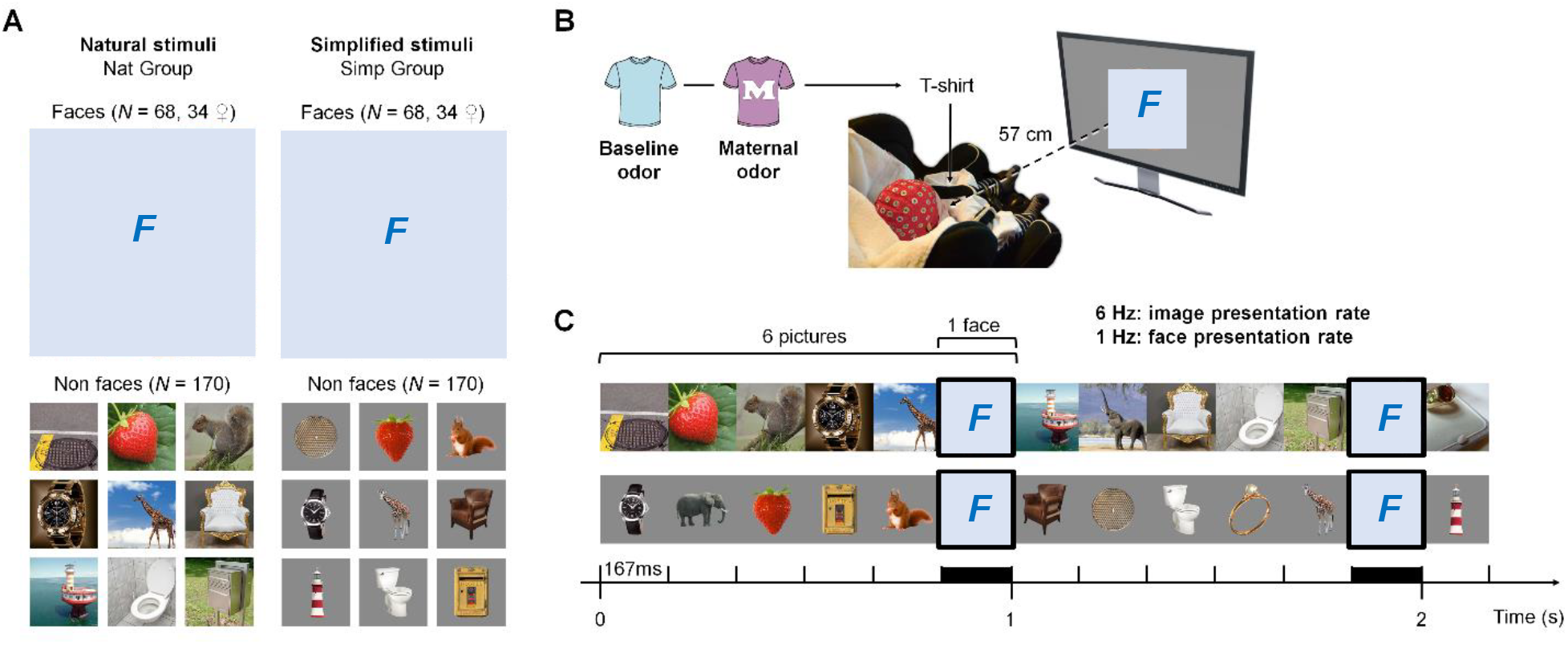
A frequency-tagging EEG approach to measure the inverse effectiveness of maternal odor on rapid face categorization. **A**. Examples of the natural (left) and simplified (right) stimuli respectively presented to Nat Group and Simp Group, each depicting human faces (*N* = 68, 34 females, here replaced by “F” placeholders for copyright issues) and nonface objects (*N* = 170, 85 categories × 2 exemplars). **B**. Two odor contexts (baseline and maternal odor) were delivered using a t-shirt disposed on the infant’s chest during visual stimulation and alternated every 2 sequences. Infants were equipped with an EEG cap and seated at 57 cm in front of the stimulation screen, in which stimuli (item + background) roughly subtended 24° of visual angle. **C**. Excerpt of 2.167 s of fast periodic visual stimulation (from 32-s-long sequences) presenting 6 images/s (i.e., at a 6-Hz rate, stimulus duration: 167 ms) with faces interspersed every 6^th^ stimulus (i.e., at a 1-Hz rate, 1-s interval between two faces).

As in previous studies showing a maternal odor effect on rapid face categorization (Leleu et al., 2020; Rekow et al., 2023, 2021, 2020), the first set (Nat Group) consisted in natural color images in which the item appears at variable locations, under variable lighting conditions, and from variable sizes and viewpoints, together with different facial expressions and external features (e.g., haircut) for faces. Items were embedded in their original background, implying figure-ground segregation. Low-level (e.g., luminance, contrast, spatial frequency) and higher-level (e.g., color, curvature) cues are thus highly variable in these natural stimuli, so that inter-stimulus variability is high both within and across face and nonface categories. This avoids the contribution of image-based characteristics to discriminate faces and nonface objects, and to generalize across individual faces, for a measure of rapid face categorization beyond physical cues (de Heering and Rossion, 2015).

The second stimulus set (Simp Group) consisted in color images depicting the exact same categories as the first set, but items were segmented from their background, which was replaced by a grey background (128/255 in greyscale). In addition, exposure conditions were less variable than in the first set (e.g., constant size, central location). In particular, faces were all depicted from a full-front view with a neutral expression under uniform lighting, and external features such as hair, neck and ears were removed. All faces covered roughly the same surface (about 200-pixel height and 150-pixel width) and were aligned at eye-level. These simplified stimuli were created to reduce physical variability across face stimuli while keeping high variability between face and nonface stimuli, thus making rapid face categorization less demanding for the visual system compared to the natural stimuli. It is also worth noting that we simplified all stimuli and not only face stimuli to avoid a face-selective response that would simply rely on low-level cues. Indeed, if we had compared simplified face stimuli to natural nonface stimuli, simple image-based characteristics would have been a strong and reliable driver of the face-selective response, although having nothing to do with face categorization (e.g., face stimuli with no background compared to nonface stimuli with a complex colorful background). Hence, even if simplified stimuli intrinsically lead to more image-based categorization, using the same type of stimuli for both face and nonface stimuli prevents from a strong contribution of low-level cues to the response. In addition, using the same manipulation for both face and nonface stimuli allows to evaluate its effect on the general visual response to all stimuli (see section 2.4 for a description of the two frequency-tagged responses).

### 2.3. Odor stimuli

Infants of both groups were exposed to two odor contexts: the mother’s body odor and a baseline odor, which were delivered using white cotton t-shirts as in previous studies (Durand et al., 2013; Leleu et al., 2020; Rekow et al., 2023, 2021, 2020). T-shirts were laundered beforehand in the lab with a scentless hypoallergenic powder detergent (Persavon, France) and then stored in a hermetic zip-locked plastic bag. The mother’s body odor was collected at home using a clean t-shirt sent to the mother one week before the experiment. Mothers were asked to wear the t-shirt on bare skin during the three consecutive nights preceding the experiment and to refrain from using odorous soap or perfume. During the days of the collection period, they were asked to store the t-shirt in the bag at room temperature, away from any heating device. The baseline odor consisted in the odor of an unworn t-shirt stored in our premises.

### 2.4. Procedure

For both groups, the procedure was identical to that of previous frequency-tagging EEG studies showing a maternal odor effect on rapid face categorization (Leleu et al., 2020; Rekow et al., 2023, 2021, 2020). After placement of the electrode cap, infants were installed in a baby car seat in front of the stimulation screen (Figure 1B) in a light- and sound-attenuated room equipped with an air-extractor located approximately 2 m above the seat and which continuously and silently renewed the air. The room was also aired between experiments and the experimenter did not use or consume any odorous product. During the experiment, infants were placed behind occluding curtains to minimize visual distraction, and continuously monitored through a camera placed on top of the screen. Parents were asked to stay at a minimal 2.5-m distance from their infant and to not interact with her/him except in case of manifest distress.

For odor delivery, the experimenter used dedicated disposable nitrile gloves (Schield Scientific, The Netherlands) to affix one t-shirt under the seat belts on the infant’s upper chest just before a sequence of visual stimulation (sequence duration: 34.5 s, see below). The t-shirt was folded to optimally expose the parts that were in contact with axillary, breast, and neck regions. Every two sequences, a break of 1 min permitted to change the t-shirt (i.e., the odor context). Both odor contexts were thus passively delivered throughout two sequences of visual stimulation, their presentation order being counterbalanced across infants.

Visual stimuli were displayed in the center of a 24-inch LED screen (refresh rate: 60 Hz, resolution: 1920 × 1080 pixels) on a grey background (128/255 in greyscale) at a viewing distance of 57 cm, thus subtending 24 × 24° of visual angle for the natural stimuli (including both the item and its background) and about half this size for the item (no visible background) in the simplified stimuli (Figure 1B). Stimuli were presented through a fast periodic visual stimulation at a rate of 6 Hz (i.e., 6 stimuli per s) without inter-stimulus interval (stimulus duration: 167 ms, i.e., 1 s/6) and faces were inserted as every 6^th^ stimulus, i.e., at a rate of 6 Hz/6^th^ = 1 Hz (1-s interval between two faces, Figure 1C). This periodic stimulation resulted in two distinct brain responses tagged at two frequencies in the EEG spectrum: a general visual response recorded at 6 Hz and harmonics (i.e., integer multiples) elicited by all visual cues rapidly changing 6 times per second (common to face and nonface stimuli), and a face-selective response recorded at 1 Hz and harmonics capturing the differential neural activity elicited by faces compared to the nonface categories and generalized across individual faces. The face-selective response is thus a signature of rapid (i.e., single-glance) face categorization (de Heering and Rossion, 2015; Jacques et al., 2016; Rossion et al., 2015).

Each sequence of visual stimulation lasted 34.5 s, starting with a pre-stimulation interval (0.5 s), followed by a fade-in of increasing contrast (0 to 100%, 1.833 s), the full-contrast stimulation (31.167 s), a fade-out of decreasing contrast (100 to 0%, 0.833 s), and a post-stimulation interval (0.167 s). For each group of infants, the set of 68 face stimuli was divided into two subsets of 34 faces (17 females) that were counterbalanced across sequences and odor contexts (all combinations reached after 4 sequences). Therefore, all face stimuli were presented equally across the two consecutive sequences of one odor context and contrasted to the same set of 170 nonface stimuli. When infants looked away from the screen, short sounds (e.g. bike ring, squeak of rubber toy) were used to reorient attention to the screen. Their sporadic and non-periodic appearances minimally contaminated the frequency-tagged EEG responses with auditory-evoked potentials. The number of sequences was not fixed a priori, such that the experiment stopped when the infant manifested disinterest or fatigue, or at parental demand.

Infants were included in the final samples if they achieved at least 4 valid sequences (i.e., 2 per odor context). A sequence was considered valid if not interrupted prematurely and if it elicited a general visual response as evidenced during preprocessing (see section 2.5). The 42 included infants achieved between 4 and 16 sequences each (mean ± *SD*: 9.4 ± 2.8 sequences), for an overall stimulation duration ranging from 2 min 18 s to 9 min 12 s per infant.

### 2.5. EEG recording and preprocessing

EEG was continuously recorded from a 63 Ag/AgCl electrode cap (Waveguard, ANT Neuro, The Netherlands) according to the 10–10 classification system (acquisition reference: CPz, ground: AFz, initial impedance < 30 kΩ, sampling rate: 1000 Hz) (Figure S2). EEG data were preprocessed and analyzed using Letswave 6 (http://www.letswave.org/) running on Matlab 2017 (MathWorks, USA).

First, we applied a Butterworth filter (highpass, cutoff: 0.1 Hz, 4^th^ order) to individual datasets. Then, the signal was down-sampled to 200 Hz before being cropped into 36-s-long segments starting from the beginning of the fade-in of each sequence. Resulting segments were cleaned of artifacts using the *Artifact Blocking* algorithm (Mourad et al., 2007) with a threshold of ± 250 μV. This algorithm removes high-amplitude artifacts without distorting other sources, thus avoiding simple artifact rejection, which cannot be applied to the long segments analyzed in frequency-tagging studies. It is particularly useful with infant EEG data, which are difficultly cleaned by other artifact-removal techniques (see Fujioka et al., 2011, for a detailed description of such difficulties and a comparison of artifact-removal techniques in infant data). Segments were then cropped again, starting from the end of the fade-in (i.e., at the onset of the first stimulus of the full-contrast phase) to the end of the fade-out, resulting in 32-s-long segments. Datasets were re-referenced according to a common average reference.

To increase signal-to-noise ratio, invalid segments were rejected using a data-driven criterion which consisted in identifying segments with no general visual response at first (6 Hz) and second (12 Hz) harmonics (used as a marker of the infant’s attention to the stimulation sequence; Leleu et al., 2020; Rekow et al., 2023, 2021, 2020). Baseline-corrected amplitude spectra were extracted for each segment and *Z*-scores were calculated (see section 2.6) to estimate significance at 4 medial occipital electrodes (POz, Oz, O1, O2) that exhibit the largest response to a 6-Hz stream of complex visual stimuli in the infant brain (e.g., Leleu et al., 2020). Segments were kept for subsequent analyses if at least two *Z*-scores were above 1.64 (*p* < .05, one-tailed, signal > noise). The total number of rejected segments was 45 out of 406 (11%). The remaining average number of segments per infant was (mean ± *SD*) 8.3 ± 2.7, and did not differ between groups (Nat Group: 9.0 ± 3.4, Simp Group: 7.7 ± 1.7, *T* (40) = 1.63, *p* = .11). When compared between odor contexts, this mean number of segments was not significantly different for both Nat Group (baseline: 4.4 ± 1.7, maternal: 4.6 ± 1.9, *T* (20) = 0.91, *p* = .37) and Simp Group (baseline: 3.8 ± 1.1, maternal: 3.9 ± 0.8, *T* (20) = 0.77, *p* = .45). Finally, remaining segments were sorted according to the odor context and averaged in the time domain to obtain a single 32-s-long segment per odor context and infant.

### 2.6. EEG frequency-domain analysis

Although the following analysis pipeline was not pre-registered, as it was not a common practice in our lab at the time we designed the study, analyses were mostly planned before data collection based on previous studies on maternal odor influence on rapid face categorization (Leleu et al., 2020; Rekow et al., 2023, 2021, 2020). First, a fast Fourier transform was applied to the two segments (one per odor context) of each infant and raw amplitude spectra were extracted for all electrodes with a frequency resolution of 1/32 s = 0.03125 Hz. Given the steep power-law function of these spectra, a baseline correction was applied to remove background noise and lead to notional amplitudes of zero in the absence of tagged responses. At each frequency bin, mean noise was estimated from 6 neighboring bins and subtracted out. These bins were selected among the 10 surrounding bins (5 on each side, ± 0.15625 Hz) after the exclusion of the 2 immediately adjacent (one on each side, in case of spectral leakage) and the 2 extreme (minimum and maximum) bins (to avoid signal and potential outliers in noise estimation). Then, we estimated the number of significant harmonics separately for each brain response using *Z*-scores calculated on the average across odor contexts, electrodes, and infants (from both groups). *Z*-scores were defined as the difference between the amplitude at the frequency of interest and the mean amplitude of 20 neighboring bins, divided by their standard deviation. Neighboring bins were selected among 22 surrounding bins beyond those used for baseline correction (11 on each side, from ± 0.1875 Hz to ± 0.5 Hz) after the exclusion of the two extreme bins. Harmonics were included until *Z*-scores were no longer consecutively significant (*Z* > 1.64, *p* < .05, one-tailed, signal > noise). For the general visual response, 4 consecutive harmonics were significant (i.e., from 6 to 24 Hz, Table S1), and 5 harmonics were significant for the face-selective response (i.e., from 1 to 5 Hz, Table S1).

Next, we identified regions of interest (ROIs) for subsequent analyses. To obtain a compiled representation of each brain response, we summed their amplitude across significant harmonics (Retter et al., 2021). Given that both responses are located over posterior brain regions (e.g., de Heering and Rossion, 2015; Leleu et al., 2020), we then explored which electrodes were significant among the 21 occipital, occipito-temporal, and parietal electrodes (Figure S2) using *Z*-scores (see above) calculated across odor contexts and infants (from both groups). *Z*-scores were considered significant when *Z* > 2.82 (*p* < .05, one-tailed, signal > noise, Bonferroni-corrected for 21 electrodes). For the general visual response, every electrode was significant (all *Z*s > 17.6, Table S2). We thus kept the 4 best electrodes, all located over the middle occipital cortex (Oz, O2, O1, POz) and identical to those of previous studies (e.g., Leleu et al., 2020; Rekow et al., 2023, 2021, 2020). For the face-selective response, 15 electrodes were significant (all *Z*s > 2.82, Table S2). We selected them (except electrode P5 due to the non-significance of its contralateral homologous electrode P6, Figure S2) to form three ROIs: right (rOT: P8, P10, PO6, PO8, O2) and left (lOT: P7, P9, PO5, PO7, O1) occipito-temporal regions, and a medial occipital region (mO: PO3, PO4, POz, Oz).

Having defined harmonics and ROIs from the mean responses across odor contexts and infants, we then conducted a two-step analysis pipeline separately on each brain response to compare (1) the two groups of infants (i.e., natural vs. simplified stimuli) irrespective of the odor context, and (2) the two odor contexts for each group of infants (i.e., each type of stimuli). For (1), we first averaged the responses across odor contexts and used *Z*-scores to determine how many harmonics were significant for each group of infants (Table S3 and Table S5 for the general and face-selective visual responses, respectively). *Z*-scores were also used to estimate the significance of the summed responses for each group and each individual infant within groups (Table S4 and Table S6 for the general and face-selective visual responses, respectively). Next, the mean amplitude of the summed brain responses was quantified for each group, and individual amplitudes were finally submitted to a repeated-measures ANOVA (Table S7) using *Group* (Nat Group, Simp Group) as a between-subject factor. For the face-selective response, *ROI* (rOT, lOT, mO) was also used as a within-subject factor and comparisons between ROIs were performed using *T*-tests. Mauchly’s test was used to estimate sphericity violation and the Greenhouse-Geisser correction for degrees of freedom (*df*) was applied whenever sphericity was violated (adjusted *df* and the epsilon coefficient (*ε*) are reported). Significance was fixed at *p* < .05 and effect sizes are reported as partial eta squared (*ηp*^2^) and/or Cohen’s *d*. To account for the potential effect of the number of averaged segments per infant (see section 2.5), we conducted additional ANCOVAs including this variable as a continuous factor together with the categorical factors used in the ANOVAs (Table S7). The main effect of this factor was significant in neither analyses and we found the same pattern of results for the other factors, except for the main effect of *ROI* on the face-selective response (see section 3.1).

For (2), the mean amplitude of the summed brain responses was first quantified for each odor context and group, and individual amplitudes were submitted to a repeated-measures ANOVA (Table S8) using the *Group* and *ROI* (for the face-selective response) factors, and adding *Odor* (baseline, maternal) as a within-subject factor. However, since in (1) we found a strong difference between groups (i.e., types of stimuli) for each brain response (i.e., larger general and face-selective visual responses to natural and simplified stimuli, respectively, see section 3.1), the *Odor* effect of the group with the weakest response might be masked by the larger response of the other group in the omnibus ANOVA. To avoid that, we normalized individual data separately for each group before running the ANOVA. We subtracted the mean amplitude across odor contexts (and ROIs for the face-selective response) and divided by its standard deviation, such that, for each group, the resulting mean amplitude and standard deviation across conditions were equal to 0 and 1, respectively (leading to a null main effect of *Group*). Given our aim to delineate the influence of maternal odor, we focused on effects involving the *Odor* factor. Significant interactions were decomposed using contrasts and paired comparisons were performed using *T*-tests. As in (1), the Greenhouse-Geisser correction was applied whenever necessary, significance was fixed at *p* < .05, effect sizes are reported as *η*_*p*_^2^ and/or Cohen’s *d*, and ANCOVAs were conducted to account for the potential effect of the number of averaged segments per infant (Table S8; no main effects of this factor, same pattern of results). To illustrate individual odor effects (maternal minus baseline) for each group, we expressed them as effect sizes by dividing by their standard deviation. Lastly, since the face-selective response to natural stimuli is mainly concentrated on the 1^st^ harmonic whereas the response to simplified stimuli is distributed on 5 harmonics (see section 3.1), we estimated whether this difference drove the odor effect by analyzing separately the 1^st^ harmonic (1 Hz) and the sum of the four remaining harmonics (2 to 5 Hz).

## 3. Results

### 3.1. General and face-selective visual responses to natural and simplified stimuli in the 4-month-old brain

As expected, a fast 6-Hz train of either natural or simplified visual stimuli elicited a large response at the same frequency in the infants’ EEG spectrum (Figure 2A). This general neural response to the rapid stimulation stream was significantly distributed until the 4^th^ harmonic (24 Hz) at medial occipital electrodes for both groups (all *Z*s > 9.73, all *p*s < .001, Table S3). The summed response across harmonics (Figure 2C) was also highly significant for both groups (Nat Group: *Z* = 113, *p* < .001; Simp Group: *Z* = 85.1, *p* < .001). Such a robust response was confirmed by individual data (Table S4) as all infants exhibited a significant general response at the medial occipital ROI (all *Z*s > 3.35, all *p*s < .001) except one in Simp Group (*Z* = 1.25, *p* = .11).

**Figure 2.**
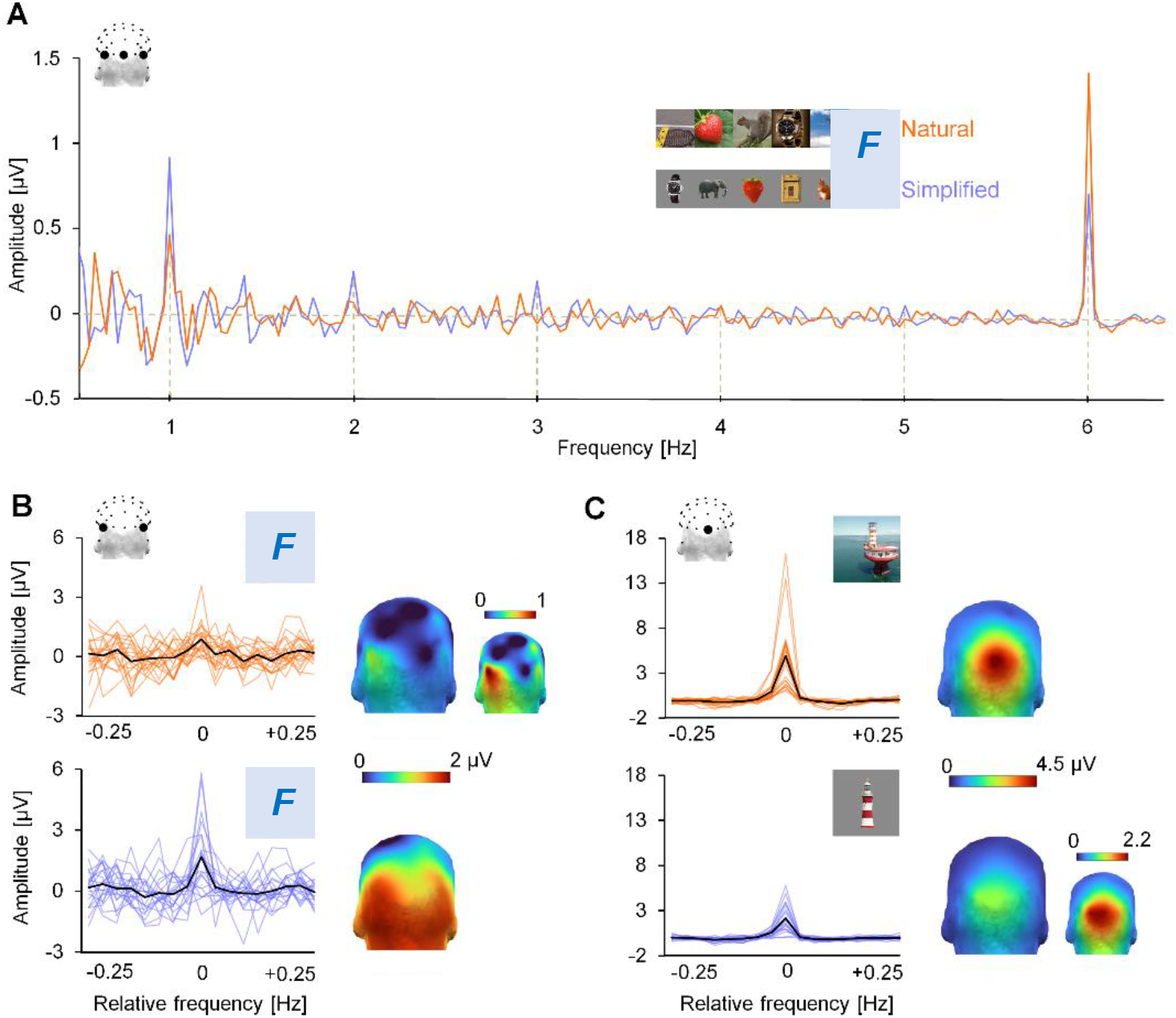
General and face-selective visual responses to natural and simplified stimuli. **A**. Mean amplitude spectrum across two occipito-temporal (PO7, PO8, best at 1 Hz) and one occipital (Oz, best at 6 Hz) electrodes averaged across odor contexts and infants for natural (orange) and simplified (purple) stimuli. The general visual response is recorded at 6 Hz and harmonics (i.e., integer multiples, not displayed) and the face-selective response at 1 Hz and harmonics (displayed from 2 to 5 Hz). **B**. and **C**. Amplitude of the face-selective (B) and general visual (C) responses (relative frequency = 0 Hz) and their surrounding noise (± 0.25 Hz) summed across 5 (face-selective) and 4 (general) harmonics and averaged across odor contexts for Nat Group (orange) and Simp Group (purple). The face-selective response is pooled across two occipito-temporal electrodes (PO7, PO8) and the general visual response is displayed at electrode Oz. The black line represents the mean of the group and colored lines represent individual responses. Topographical head maps (posterior view) illustrate the spatial distribution of the responses. Smaller maps illustrate the lowest responses with an adjusted scale (i.e., face-selective response to natural stimuli in B, general visual response to simplified stimuli in C). Face stimuli are replaced by “F” placeholders for copyright issues.

Interestingly, however, visual inspection also indicated a stronger general visual response to natural than simplified stimuli (Figure 2). At electrode Oz where the highest response was recorded for both types of stimuli, amplitude was 123% larger (Cohen’s *d* = 1.01) for Nat Group (4.90 ± 0.84 (*SEM*) μV) than for Simp Group (2.20 ± 0.33 μV). This difference was smaller but remained strong (+89%, Cohen’s *d* = 0.87) for the mean amplitude across medial occipital electrodes (Nat Group: 3.32 ± 0.49 μV, Simp Group: 1.76 ± 0.30 μV) and was statistically significant, as revealed by the main effect of *Group* (*F* (1, 40) = 7.47, *p* = .009, *η*_*p*_^2^ = .16).

Another neural response was visible in the EEG spectrum at the 1-Hz frequency of face presentation and its harmonics (Figure 2A). Contrary to the general visual response, the face-selective response appeared especially large and distributed on several harmonics for Simp Group, while lower and mainly visible at the 1^st^ harmonic (1 Hz) for Nat Group. The response to simplified stimuli was indeed significant until the 5^th^ harmonic (5 Hz) at almost every ROI and for the mean across ROIs (Table S5), with *Z*-scores ranging between *Z* = 2.64 (*p* = .004, 4^th^ harmonic) and *Z* = 9.24 (*p* < .001, 1^st^ harmonic). Summed across harmonics (Figure 2B), it was significant at every ROI and for the mean across ROIs (all *Z*s > 8.77, all *p*s < .001). In contrast, the face-selective response to natural stimuli was almost exclusively significant at the left-hemispheric ROI (lOT) for the 1^st^ and 4^th^ harmonics, which were also the only significant harmonics for the mean across ROIs (1^st^ harmonic: *Z* = 2.09, *p* = .018; 4^th^ harmonic: *Z* = 2.19, *p* = .014; other harmonics: all *Z*s < 1.20, all *p*s > .11). The summed response reached significance only in the left hemisphere (lOT) and for the mean across ROIs (all *Z*s > 2.37, all *p*s < .009), with a marginal response in the right hemisphere (rOT: *Z* = 1.38, *p* = .084) (Table S5). Despite this difference between groups, individual data (Table S6) indicated that the face-selective response was driven by a large subset of infants in both groups, 20 infants out of 21 (95%) showing at least one significant electrode within the ROIs for Simp Group, and 16 out of 21 (76%) for Nat Group.

Direct comparison between groups confirmed that the mean face-selective response across ROIs was significantly much stronger (+350%, Cohen’s *d* = 1.29) for simplified than for natural stimuli, with an amplitude of 1.50 ± 0.28 μV for Simp Group as opposed to 0.33 ± 0.11 μV for Nat Group (main effect of *Group*: *F* (1, 40) = 14.9, *p* < .001, *η*_*p*_^2^ = .27). There was also a significant main effect of *ROI* (*F* (1.5, 61.5) = 3.91, *ε* = 0.77, *p* = .035, *η*_*p*_^2^ = .09) that did not interact with *Group* (*F* < 1) and did not survive after including the number of averaged segments per infant in the analyses (*F* < 1, Table S7). Nevertheless, the face-selective response was particularly large at lOT in Nat Group, as lOT (0.58 ± 0.20 μV) represented 58% of the overall response across ROIs *vs*. 28% for rOT (0.28 ± 0.12 μV) and only 14% for mO (0.14 ± 0.15 μV). The topography of the response was more balanced in Simp Group with 38% for lOT (1.71 ± 0.30 μV), 34% for rOT (1.52 ± 0.37 μV), and 28% for mO (1.26 ± 0.27 μV).

In sum, a 6-Hz train of natural visual stimuli triggered a larger medal occipital response than a train of simplified stimuli, whereas the appearance of human faces at 1 Hz within such rapid stimulations elicited a weaker occipito-temporal face-selective response to natural than simplified face stimuli, reflecting more demanding visual categorization for the 4-month-old brain.

### 3.2. Maternal odor selectively improves the more demanding categorization of natural face stimuli

Visual inspection of the face-selective response dissociated between groups and odor contexts suggested that the mother’s body odor enhanced the response to natural stimuli over the right occipito-temporal cortex while the response to simplified stimuli was less affected in any region (Figure 3A). We indeed found a significant *Odor* × *Group* interaction (*F* (1, 40) = 5.19, *p* = .028, *η*_*p*_^2^ = .11) due to a significant *Odor* effect for natural (*F* (1, 40) = 7.66, *p* = .009, *η*_*p*_^2^ = .16) but not simplified (*F* < 1) visual stimuli. There was also a significant interaction between the *Odor* and *ROI* factors (*F* (2, 80) = 3.14, *p* = .049, *η*_*p*_^2^ = .07) revealing an *Odor* effect only at rOT (*F* (1, 40) = 9.02, *p* = .005, *η*_*p*_^2^ = .18; two other regions: both *F*s < 1).

**Figure 3.**
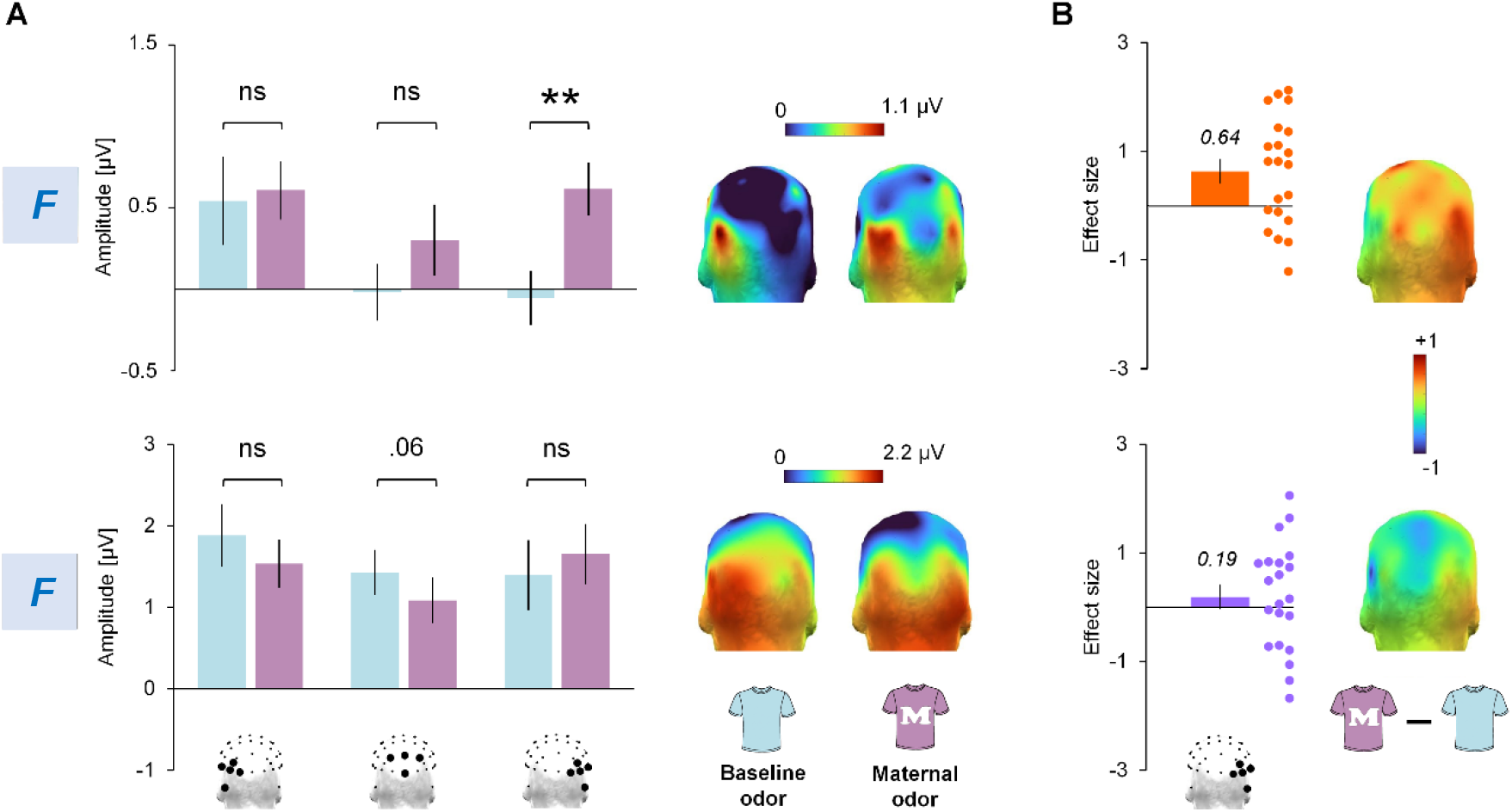
Face-selective response to each type of stimuli (group of infants) as a function of the odor context. **A**. Amplitude of the face-selective response at the left occipito-temporal (lOT: P7, P9, PO5, PO7, O1), medial occipital (mO: POz, PO3, Oz, PO4), and right occipito-temporal (rOT: P8, P10, PO6, PO8, O2) regions for natural (top) and simplified (bottom) stimuli in the baseline (blue) and maternal (violet) odor contexts. Error bars represent standard errors of the mean (** *p* = .008, ns *p* >. 25, *p* = .06 for mO and simplified stimuli). Topographical head maps (posterior view) illustrate the spatial distribution of the response. **B**. Mean (box) and individual (circles) effect sizes (Cohen’s *d*s) of the odor effect (maternal – baseline) at rOT. Error bars represent standard errors of the mean and head maps (posterior view) illustrate the topography of the effect size. Face stimuli are replaced by “F” placeholders for copyright issues.

Comparison between odor contexts for each group and each ROI confirmed that the face-selective response to natural stimuli recorded at the right occipito-temporal region (rOT) was significantly larger in the maternal (0.62 ± 0.16 μV) than in the baseline (−00.05 ± 0.16 μV) odor context (*T* (20) = 2.95, *p* = .008, Cohen’s *d* = 0.64). In contrast, the effect of the mother’s body odor was non-significant at mO (maternal: 0.30 ± 0.22 μV, baseline: −0.02 ± 0.17 μV, *T* (20) = 1.16, *p* = .26, Cohen’s *d* = 0.25) and lOT (maternal: 0.61 ± 0.18 μV, baseline: 0.54 ± 0.27 μV, *T* < 1, Cohen’s *d* = 0.06). Interestingly, the face-selective response was entirely distributed at lOT in the baseline odor context, whereas the response was spatially more extended in the maternal odor context with 40%, 40%, and 20% of the overall response across ROIs at lOT, rOT, and mO, respectively.

For simplified stimuli, the face-selective response slightly increased (+19%) in the mother’s odor context over the right hemisphere (rOT: maternal: 1.66 ± 0.37 μV, baseline: 1.39 ± 0.43 μV) and decreased (−018%) over the left hemisphere (lOT: maternal: 1.54 ± 0.29 μV, baseline: 1.88 ± 0.38 μV), but these effects failed to reach significance (both *T*s < 1.05, both *p*s > .30, both *d*s < .23). Though stronger over the middle occipital cortex (mO), the diminution of the response (−024%) with maternal odor (maternal: 1.08 ± 0.28 μV, baseline: 1.43 ± 0.28 μV) was only a trend (*T* (20) = 1.99, *p* = .061, Cohen’s *d* = 0.43). As a result, contrary to Nat Group, the topographical distribution of the response across ROIs in Simp Group remained quite stable between odor contexts, with 40% at lOT, 30% at rOT, and 30% at mO in the baseline odor context as opposed to 36% at lOT, 39% at rOT, and 25% at mO in the maternal odor context. Mean and individual odor effects at rOT in both groups of infants are depicted in Figure 3B as effect sizes (i.e., Cohen’s *d*s, medium effect for Nat Group, *d* = 0.64, negligible effect for Simp Group, *d* = 0.19).

In sum, while the face-selective response to simplified stimuli did not significantly change with maternal odor, the response to natural face stimuli evolved from a focal activity at left occipito-temporal sites in the baseline odor context to a more widespread response, especially in the right hemisphere, when adding the mother’s body odor.

Finally, given that the face-selective response to natural stimuli was mainly concentrated on the 1^st^ harmonic (1 Hz) whereas the response to simplified stimuli was distributed on 5 harmonics (from 1 to 5 Hz), we determined whether the odor effect observed at rOT in Nat Group depended on these different frequency representations of the response. When considering only the 1^st^ harmonic (1 Hz, Figure 4A), the amplitude of the face-selective response to natural stimuli was still significantly larger in the maternal (0.48 ± 0.15 μV) than in the baseline (−00.04 ± 0.13 μV) odor context (*T* (20) = 2.38, *p* = .027, Cohen’s *d* = 0.52). In contrast, for the summed response across remaining harmonics (from 2 to 5 Hz, Figure 4B), there was no significant difference between the mother’s (0.13 ± 0.12 μV) and the baseline (−00.02 ± 0.09 μV) odors (*T* (20) = 0.92, *p* = .37, Cohen’s *d* = 0.20). Likewise, the odor effect on the face-selective response to simplified stimuli was neither significant at 1 Hz (maternal: 1.01 ± 0.29 μV, baseline: 0.84 ± 0.26 μV, *T* (20) = 0.68, *p* = .51, Cohen’s *d* = 0.15), nor for the sum from 2 to 5 Hz (maternal: 0.64 ± 0.14 μV, baseline: 0.55 ± 0.22 μV, *T* (20) = 0.48, *p* = .63, Cohen’s *d* = 0.11). Hence, the odor effect on the face-selective response to natural stimuli was mainly driven by the 1^st^ harmonic, and no odor effect was masked by harmonic summation for the response to simplified stimuli.

**Figure 4.**
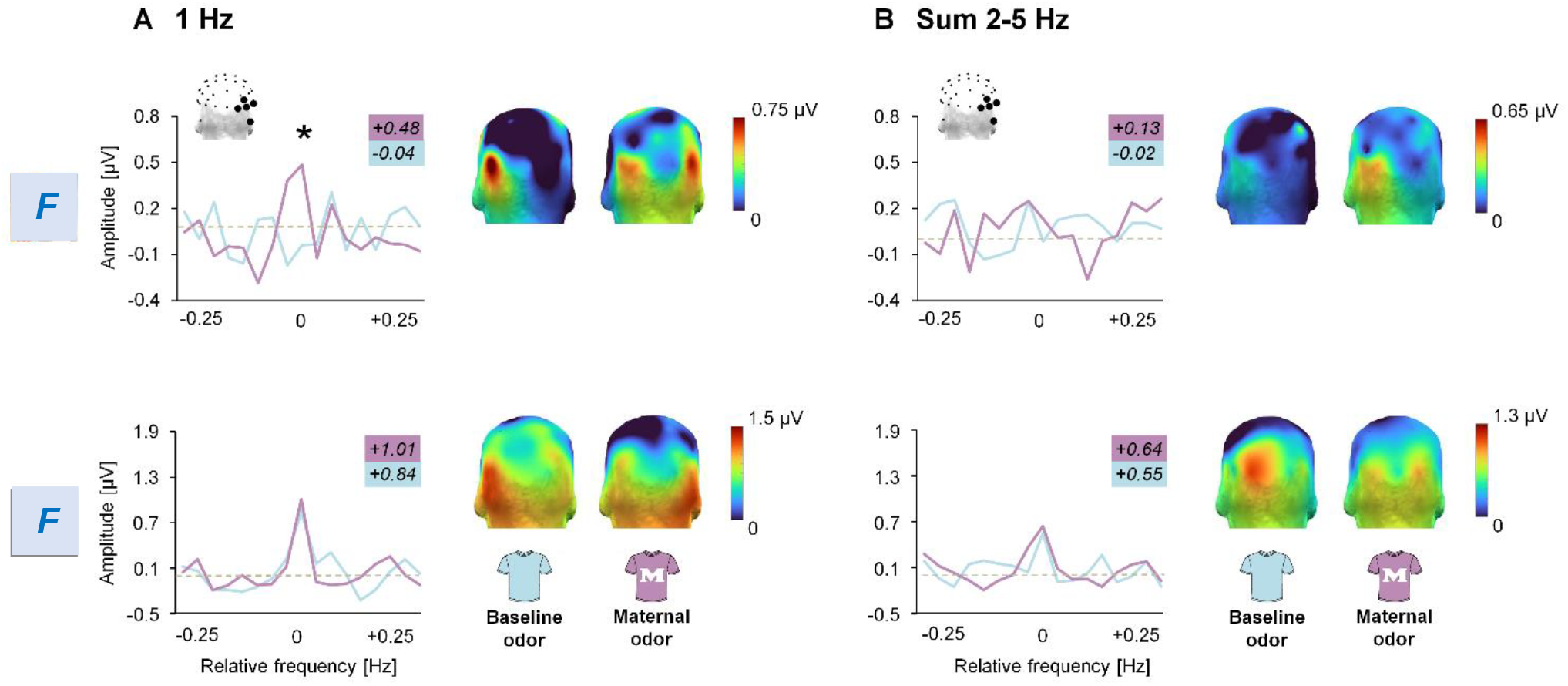
First and other harmonics of the face-selective response to each type of stimuli (group of infants) as a function of the odor context. Amplitude of the face-selective response (relative frequency = 0 Hz) and its surrounding noise (± 0.25 Hz) for the 1^st^ (1 Hz, **A**) and for the sum from the 2^nd^ to 5^th^ (2-5 Hz, **B**) harmonics at rOT for natural (top) and simplified (bottom) stimuli in the baseline (blue) and maternal (violet) odor contexts. The dashed grey line represents 0 μV. Topographical head maps (posterior view) illustrate the spatial distribution of the response. A significant difference between the two odor contexts was found at 1 Hz for natural stimuli (* *p* = .027). Face stimuli are replaced by “F” placeholders for copyright issues.

### 3.3. No odor effect on the general visual response to either type of stimuli

As noticeable in Figure 5, the general visual response to a fast train of natural stimuli was slightly larger with maternal odor while the opposite was apparent for the response to simplified stimuli. However, neither the main effect of *Odor* (*F* < 1) nor the *Odor* × *Group* interaction (*F* (1, 40) = 1.78, *p* = .19, *η*_*p*_^2^ = .04) reached significance. For natural stimuli, the amplitude of the general visual response did not significantly increase in the maternal (3.51 ± 0.58 μV) compared to baseline (3.13 ± 0.51 μV) odor context (*T* (20) = 0.80, *p* = .43, Cohen’s *d* = 0.18), and for simplified stimuli, it did not significantly decrease in the maternal (1.63 ± 0.29 μV) compared to baseline (1.89 ± 0.34 μV) odor context (*T* (20) = 1.13, *p* = .27, Cohen’s *d* = 0.25). Therefore, despite descriptive trends, the general visual response to either type of stimuli remained immune to the presence of the mother’s odor.

**Figure 5.**
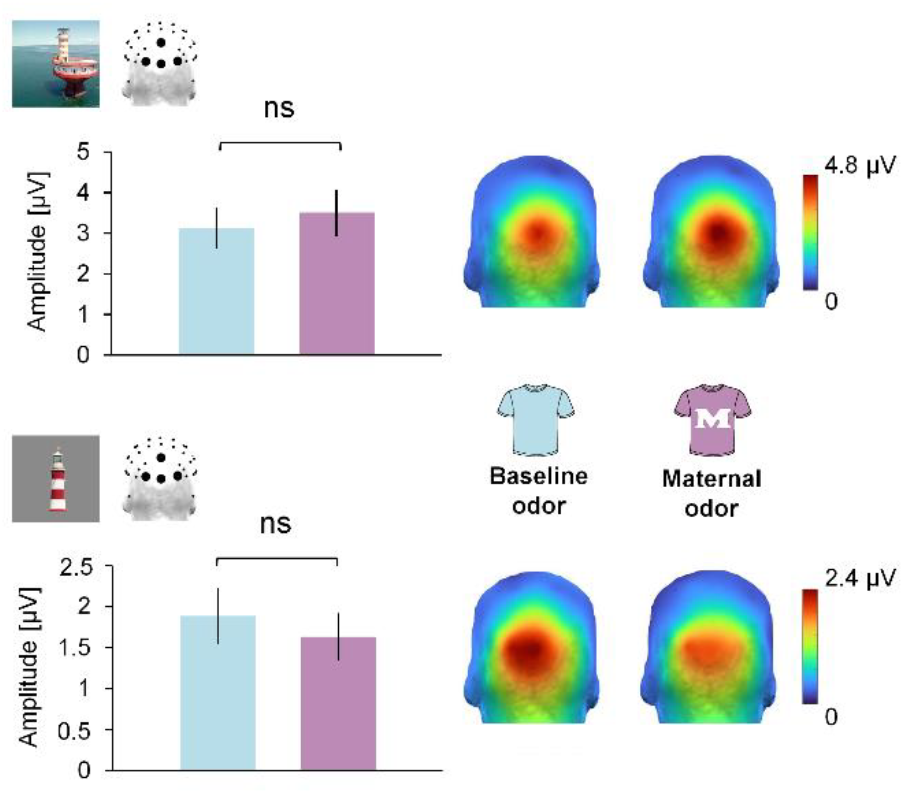
General visual response to each type of stimuli (group of infants) as a function of the odor context. Amplitude of the general visual response at the medial occipital region (POz, O1, Oz, O2) for natural (top) and simplified (bottom) stimuli in the baseline (blue) and maternal (violet) odor contexts. Error bars represent standard errors of the mean (ns *p* > .26). Topographical head maps (posterior view) show the spatial distribution of the response.

## 4. Discussion

By using frequency-tagging EEG to measure rapid face categorization in two groups of 4-month-old infants, and by exposing them to their mother’s *vs*. a baseline odor, the present study provides two major findings. First, the ability to rapidly (i.e., at a glance) categorize a variety of human faces at 4 months depends on the complexity of visual stimuli, improving when they are simplified compared to natural stimuli. Second, the mother’s body odor fosters rapid face categorization only when perceptual demand is high (natural stimuli), this effect disappearing when perceptual demand is reduced (simplified stimuli). Therefore, our results provide the first evidence that olfactory-to-visual facilitation follows the inverse effectiveness principle in the 4-month-old brain. They extend previous audiovisual studies in infants (e.g., Bahrick et al., 2010) and generalize them to olfactory-visual displays.

The use of two types of visual stimuli in a frequency-tagging design revealed a clear dissociation between the two tagged brain responses. The general visual response, elicited by the fast 6-Hz stimulation stream, is a large and reliable medial occipital activity recorded for both natural and simplified stimuli. However, it is significantly two times larger for natural than for simplified stimuli. This neural activity reflects the mere response to all cues that change 6 times per second, including both low-level (e.g., contrasts, spatial frequencies) and higher-level (e.g., colors) cues (Norcia et al., 2015). Hence, since natural stimuli depict items and their background under variable exposure conditions, these cues are highly heterogenous and trigger a larger general visual response than the lower physical heterogeneity across simplified stimuli, which depict only items (no background) under less variable conditions.

In contrast, the periodic appearance of human faces at 1 Hz within the train of stimuli elicits a clearly (4.5 times) weaker occipito-temporal response to natural than to simplified face images, which is mainly left-hemispheric and concentrated on the first harmonic for natural stimuli, while more extensively distributed over posterior brain regions and significant for five harmonics for simplified stimuli^1^. This face-selective response is a signature of rapid face categorization in the brain – a differential neural activity elicited by the reliable discrimination between the various faces and nonface objects displayed in a sequence (de Heering and Rossion, 2015; Jacques et al., 2016; Rossion et al., 2015). Previous studies using the same approach found a weak and single-harmonic response to natural face stimuli at 4 months (e.g., de Heering and Rossion, 2015; Leleu et al., 2020). Later on, the response to natural faces increases and becomes distributed on more harmonics in older infants (Rekow et al., 2023), children (Lochy et al., 2019; Vettori et al., 2019), and adults (Jacques et al., 2016; Rossion et al., 2015), reflecting improved face categorization with maturation and development. Similarly, here, the larger and more complex response to simplified stimuli indicates that edited pictures of faces, all displayed with similar size, viewpoint, expression, background, etc., are more readily categorized than widely variable natural face stimuli. As intended, our simplified stimuli reduced perceptual demand for the 4-month-old brain. In particular, while natural stimuli imply categorization beyond mere physical cues (de Heering and Rossion, 2015), image-based characteristics (e.g., local contrasts) may contribute more to the response to simplified stimuli, as suggested by a greater contribution of the medial occipital region (+14%) compared to natural stimuli.

When dissociating the two odor contexts, the general response to either natural or simplified stimuli remains of similar magnitude, confirming prior evidence that the mother’s body odor does not have a mere undifferentiated impact on visual attention (e.g., Leleu et al., 2020). In contrast, the face-selective response to natural stimuli is only recorded at left occipito-temporal sites in the baseline odor context, and increases in the right hemisphere in the presence of the mother’s body odor. This result replicates previous studies using the same EEG approach in 4-month-old infants, documenting larger right-hemispheric face-selective activity during exposure to maternal odor, either with human faces (Leleu et al., 2020; Rekow et al., 2023) or face-like objects (Rekow et al., 2021). Although future studies should delineate which odor cues drive the effect, and whether other body odors can foster face categorization in young infants, as found in adults (e.g., Rekow et al., 2022), this reinforces accumulating evidence that the mother’s odor influences face perception in infancy (Durand et al., 2020, 2013; Endevelt-Shapira et al., 2021; Jessen, 2020).

The mother’s body odor is a potent stimulus that promotes a variety of responses from the very beginning of life and often accompanies infants during their first social interactions as they are often carried by their mother while being exposed to various faces (Schaal et al., 2020 for review). This creates optimal conditions for an association from repeated co-exposure to maternal odor and (visual) faces, leading to intersensory facilitation, as previously found for faces and voices (e.g., Hyde et al., 2011). At the brain level, this could shape reentrant connectivity between the olfactory system and face-selective regions in the ventral visual stream (Edelman, 1993), as suggested by adult studies showing that the lateral fusiform gyrus, which is part of the face-selective network, responds to body odors (e.g., Zhou and Chen, 2008) and interacts with the primary olfactory cortex (Zhou et al., 2019). Thanks to this connectivity, (maternal) body odors would become able to augment the sensitivity of face-selective regions, which in turn would more effectively respond to face(−0like) stimuli in the environment. Moreover, the greater contribution of the right hemisphere, dominant for face categorization (Rossion and Lochy, 2022, for review), and more engaged in odor recognition (Brand et al., 2001; Royet, 2004), suggests that the mother’s odor triggers a “high-level” response to faces in the right occipito-temporal cortex. This interpretation is also suggested by a trend for a reduced contribution of the medial occipital region when simplified face stimuli are perceived in the maternal odor context.

However, and importantly, no enhancement of the face-selective response to simplified stimuli was found with maternal odor. This result remained unchanged when the first harmonic was analyzed separately, ensuring that the effect was not masked by the compiled representation of the response across harmonics. Therefore, according to our main hypothesis, olfactory-to-visual facilitation is found only when perceptual demand is high for the 4-month-old brain, i.e., when faces must be rapidly categorized from variable natural stimuli. This extends to olfaction and vision previous audiovisual studies showing that intersensory facilitation depends on unisensory perceptual demand at a given age (e.g., Bahrick et al., 2010). More generally, this agrees with the view that intersensory facilitation is mainly found in early development when unisensory perception is not yet fully effective by itself, and gradually attenuates as unisensory perception develops and improves (Bahrick and Lickliter, 2012), as characterized in the audiovisual domain (e.g., Bahrick and Lickliter, 2004) and recently for olfactory-visual interactions (Rekow et al., 2023).

This relative weight of intersensory facilitation along development aligns with the inverse effectiveness principle, whereby the strength of multisensory integration is a function of the effectiveness of unisensory perception (e.g., Meredith and Stein, 1983; Stevenson et al., 2007). Multisensory cues help indeed to disambiguate difficult-to-perceive unisensory inputs (Ernst and Bülthoff, 2004) and ambiguous, degraded or noisy stimuli are accordingly used to demonstrate inverse effectiveness in adults (e.g., Regenbogen et al., 2016; Stevenson et al., 2012). Similarly, for olfactory-visual interactions in adults, a body odor facilitates the rapid categorization of ambiguous face-like objects whereas the categorization of human faces, which is already effective from the sole visual input, remains immune to the odor effect (Rekow et al., 2022). Hence, in lowering visual demand, the present study indicates that inverse effectiveness already applies to olfactory-to-visual facilitation in the 4-month-old brain. Future studies should pursue this investigation in rendering face categorization more demanding at an age when no odor effect is observed under similar viewing conditions (e.g., 12 months; Rekow et al., 2023), to determine whether an olfactory-to-visual facilitation reappears.

Admittedly, the present study suffers from some limitations. As mentioned above, we do not know which odor cues in the mother’s odor actively influence face categorization, and one could even wonder whether any odor elicits the same effect. We used the mother’s body odor for its effectiveness on infant behavior and cognition (e.g., Durand et al., 2020, 2013; Endevelt-Shapira et al., 2021; Jessen, 2020), especially its selective influence on the rapid categorization of faces, either genuine (Leleu et al., 2020; Rekow et al., 2023) or illusory (Rekow et al., 2021), as opposed to nonface categories (e.g., cars; Rekow et al., 2020). Moreover, given that (1) a generic body odor (i.e., a composite odor pooled from 8 donors) selectively facilitates the rapid categorization of face-like objects in adults, contrary to a non-body (gasoline) odor (Rekow et al., 2022), (2) congruency effects between body odors and face perception beyond face categorization were repeatedly reported at various ages (Damon et al., 2021, for review), and (3) olfactory-visual interactions beyond body odors and face perception in adults imply a congruent association between odor and visual stimuli (e.g., Lundström et al., 2019; Rekow et al., 2022; Seigneuric et al., 2010; Zhou et al., 2010), we assumed that the mother’s odor is (one of) the most congruent odors for face perception in young infants. As previously mentioned, this congruency would simply emerge from the repeated co-occurrence of the mother’s odor and faces in the infant’s environment, in line with a broadly tuned perceptual system that assembles events from their simultaneity (Murray et al., 2016). Later on, this association may progressively shift as perceptual abilities develop and the environment changes (see Rekow et al., 2023, for a discussion). Whether other odors may influence face categorization in the infant brain, and whether the relation between maternal odor and (visual) faces may progressively change with age, await further assessment.

It is also worth noting that the selective response to natural face stimuli is surprisingly left-lateralized in the baseline odor context whereas previous studies using the same stimuli found a significant response over the right hemisphere (Leleu et al., 2020; Rekow et al., 2023). One reason for this discrepancy could come from mask use during the Covid-19 pandemic. Data collection indeed occurred between March 2021 and February 2022, when people were often wearing masks, and recent studies indicate that mask use had an effect on face perception in infants (Galusca et al., 2023) and children (Stajduhar et al., 2022; but see Yates and Lewkowicz, 2023). In making infants focus on less features only in the upper part of the face, masks may have favored a left-hemispheric response to faces compared to full face perception. Another non-mutually exclusive interpretation could be that the current and previous studies were underpowered, thus producing inconsistent effects. However, despite a slightly lower effect size (Cohen’s *d* = 0.64) compared to the one used for sample size estimation (Leleu et al., 2020; Cohen’s *d* = 0.79), the observed power is 83%, which remains acceptable given the conventional power level of 80% that is often unreached in infant studies (see e.g., Oakes, 2017, for a review of 70 behavioral studies published between 2013 and 2015). Although future studies should elaborate on this issue^2^, we nevertheless replicated the maternal odor effect at right occipito-temporal scalp locations (Leleu et al., 2020; Rekow et al., 2023, 2021, 2020), suggesting that the mother’s odor solicits the dominant right hemisphere for face categorization in any case.

In conclusion, we have evidenced the inverse effectiveness of olfactory-to-visual facilitation in the 4-month-old brain using a well-established maternal odor effect on rapid face categorization and making categorization less demanding with simplified stimuli. This indicates that the principles subtending how the senses interact apply to olfaction despite functional dissimilarities with the other senses, such as its reduced spatiotemporal sensitivity compared to vision and audition (Sela and Sobel, 2010). This means that contrary to audiovisual perception, which relies on a precise spatiotemporal synchrony in infants (Lewkowicz, 1996; Neil et al., 2006; Werchan et al., 2018), olfactory-visual interactions operate on a broader spatiotemporal relation that does not hamper their association, and may even favor categorization by increasing generalization across variable signals. This should be further investigated by comparing the ability of odors and sounds to improve visual categorization. In sum, our findings indicate that olfaction participates in the concert of the senses from early on in helping young infants to navigate their complex environment when unisensory perception is not fully effective by itself.

## Endnotes

1 Harmonics in the frequency domain represent the complexity (i.e., nonlinearity) of the response in the time domain. A response with several short components in time is represented by several harmonics in the frequency spectrum, whereas a single and slow response is represented by less harmonics (Retter et al., 2021).

2 To estimate an adequate sample size for future studies, we determined what might be the true effect size of maternal odor on frequency-tagged face-selective neural activity at 4 months by considering the present data (*d* = 0.64, *N* = 21), the data of Leleu et al. (2020; *d* = 0.79, *N* = 18), and the data of Rekow et al. (2023; first half of the sample aged from 4 to 8 months, *d* = 0.82, *N* = 25). This yielded a weighted mean effect size of 0.75 (95% confidence interval: 0.50–1.00). Sample size would thus be *N* = 14 with a conventional power level of 80%, and *N* = 23 with a more stringent level of 95%.

## Data availability

Anonymized preprocessed EEG data and participant information will be made permanently and publicly available once the paper is accepted for publication. They can be made privately accessible upon request to the corresponding author.

## Ethics approval

Testing was approved by the French ethics committee (CPP Sud-Est III - 2016-A02056-45).

## Acknowledgments

The authors are grateful to the participating parents and infants. They also wish to thank Véronique Boulanger for her help in recruiting them.

