## Supplement for "Olfactory facilitation of visual categorization in the 4-month-old brain depends on visual demand"

### *Supporting Information*

A

B

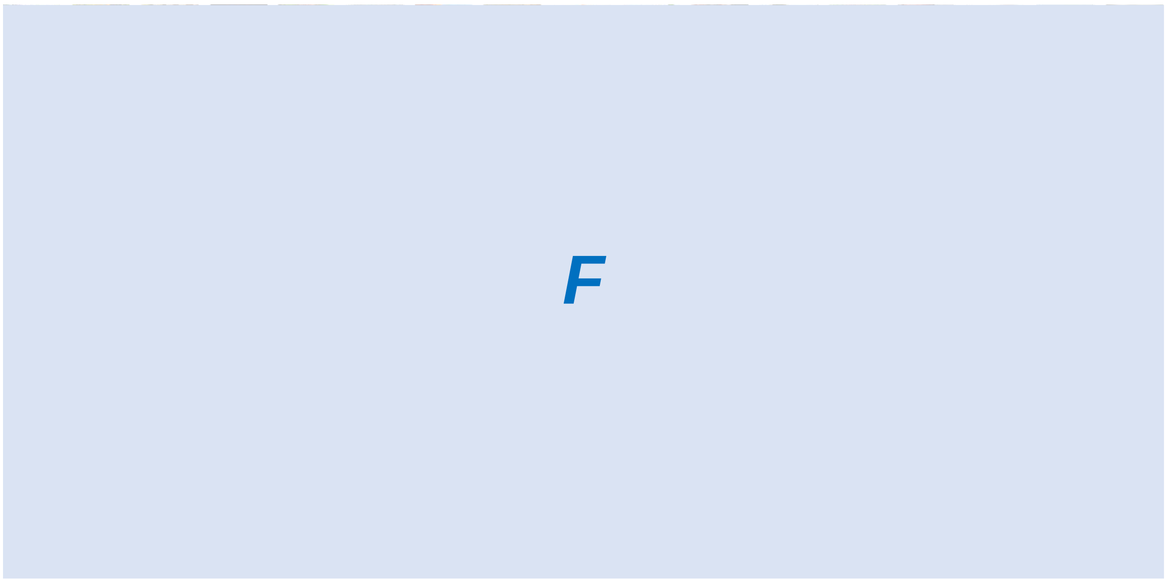

C

D

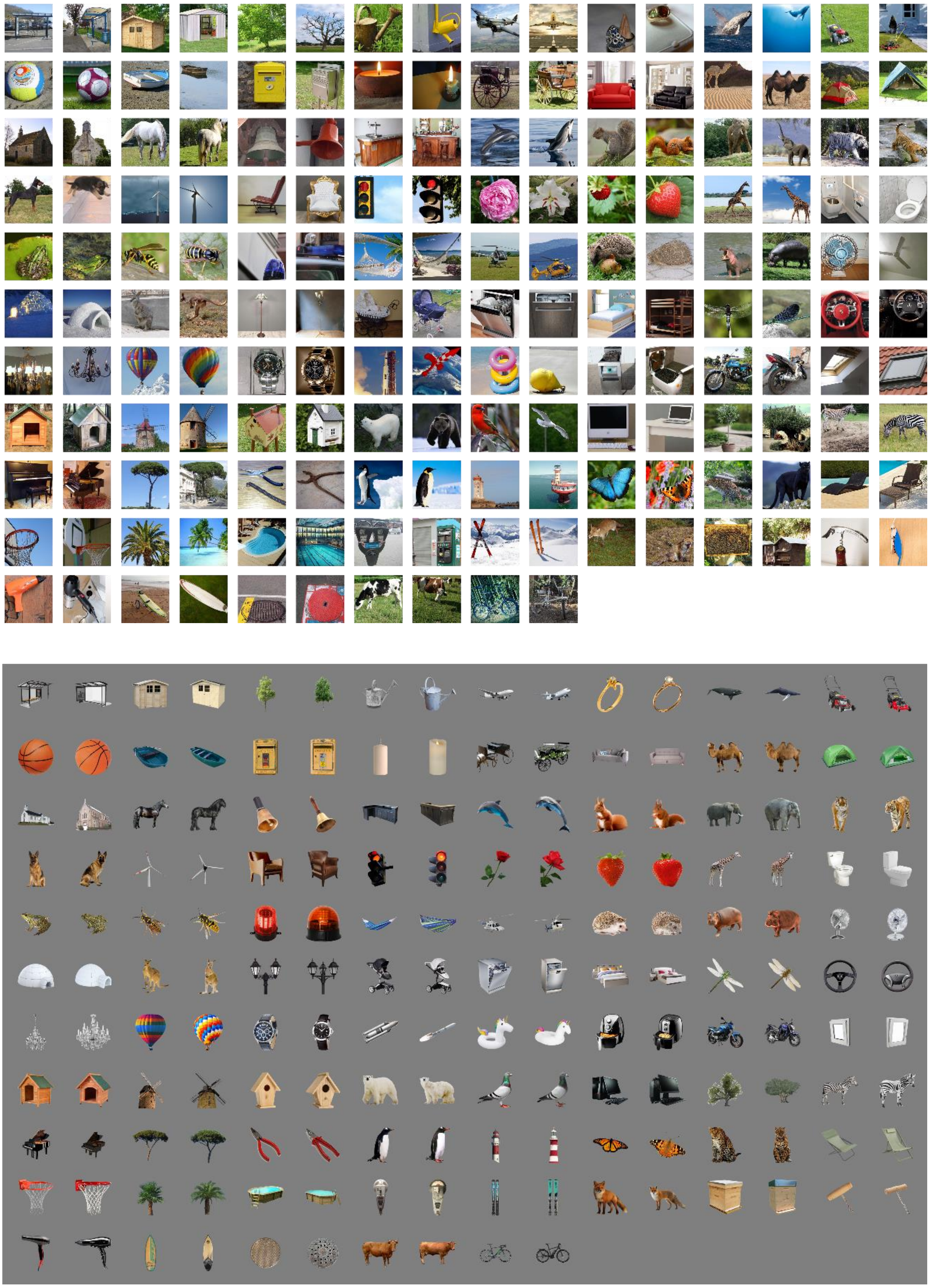

**Figure S1. Natural (Nat Group; A and C) and simplified (Simp Group; B and D) stimuli.** The two visual stimulus sets comprised pictures of 68 faces (34 females; A and B) and 170 living and non-living objects (85 categories × 2 exemplars; C and D). Face stimuli are replaced by "F" placeholders for copyright issues.

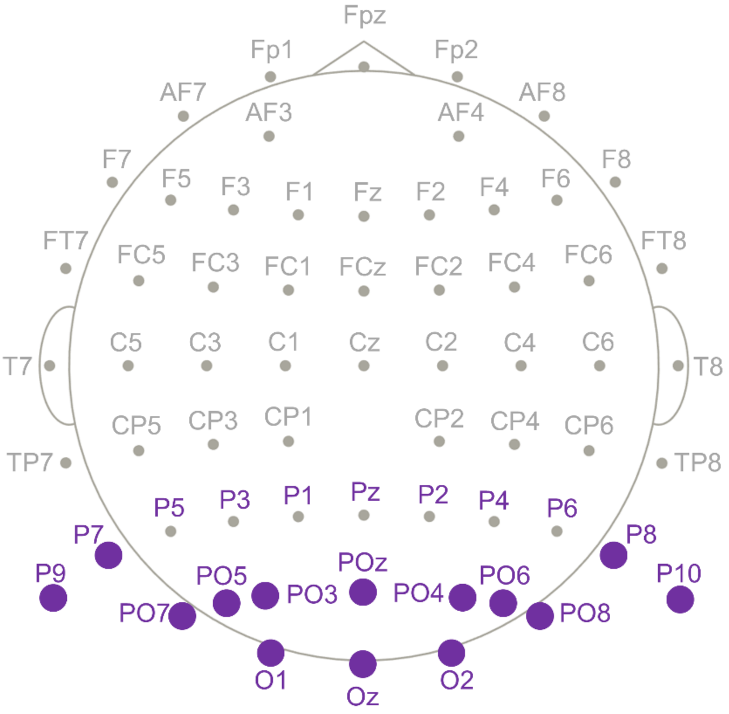


**Figure S2. Sixty-three-channel montage of EEG acquisition and electrodes considered for analysis**. EEG was acquired from a 63-channel head-cap configured according to the 10-10 classification system. Frequency-domain analysis was conducted on 21 posterior electrodes (labels colored in purple). For the face-selective response, 14 electrodes (identified by purple disks) were selected to define regions of interest (ROIs).

**Table S1. Harmonic significance for the general and face-selective visual responses.** For each response, harmonic significance was estimated using $Z$-scores calculated on the average of all electrodes, odor contexts and infants. Harmonics were considered significant according to a threshold of $Z > 1.64$ ($p < .05$, one-tailed, signal > noise). For the face-selective response, the 6th harmonic was not considered as it corresponds to the 1st harmonic of the general visual response. Asterisks indicate significance (* $p < .05$, ** $p < .01$, *** $p < .001$).

| | **General visual response** | | **Face-selective response** | |
|---|---|---|---|---|
| **Harmonic** | **Frequency (Hz)** | ***Z*-score** | **Frequency (Hz)** | ***Z*-score** |
| 1 | 6 | 70.5*** | 1 | 7.07*** |
| 2 | 12 | 71.2*** | 2 | 3.66*** |
| 3 | 18 | 37.4*** | 3 | 5.91*** |
| 4 | 24 | 11.7*** | 4 | 2.60** |
| 5 | 30 | 1.06 | 5 | 1.65* |
| 6 | 36 | -0.35 | | |
| 7 | 42 | 0.25 | 7 | 1.27 |
| 8 | 48 | -0.84 | 8 | 0.71 |

**Table S2. Electrode significance for the general and face-selective visual responses.** For each response, electrode significance was estimated for 21 posterior electrodes (Figure S2) using *Z*-scores calculated on responses summed across significant harmonics and averaged across odor contexts and infants. Electrodes were considered significant according to a threshold of $Z > 2.82$ ($p < .05$, one-tailed, signal > noise, Bonferroni-corrected for 21 electrodes). *Z*-scores are presented in decreasing order and asterisks indicate significance (* $p < .05$, *** $p < .001$). Electrodes highlighted in color were used to define ROIs (grey: medial occipital, blue: left occipito-temporal, orange: right occipito-temporal).

| | General visual response | | Face-selective response | |
|---|---|---|---|---|
| **Rank** | **Electrode** | ***Z*-score** | **Electrode** | ***Z*-score** |
| 1 | Oz | 106*** | P9 | 9.05*** |
| 2 | O2 | 101*** | PO7 | 8.69*** |
| 3 | O1 | 92.2*** | P10 | 6.95*** |
| 4 | POz | 73.2*** | PO8 | 6.56*** |
| 5 | PO3 | 68.5*** | O1 | 6.52*** |
| 6 | PO5 | 68.0*** | PO4 | 5.34*** |
| 7 | PO4 | 63.4*** | P7 | 5.20*** |
| 8 | PO6 | 59.9*** | PO5 | 5.06*** |
| 9 | PO7 | 41.2*** | P5 | 4.50*** |
| 10 | PO8 | 40.6*** | P8 | 4.05*** |
| 11 | P8 | 37.8*** | PO6 | 3.99*** |
| 12 | P6 | 32.1*** | Oz | 3.13* |
| 13 | P3 | 26.9*** | PO3 | 3.12* |
| 14 | P1 | 25.1*** | POz | 3.05* |
| 15 | P5 | 24.4*** | O2 | 2.83* |
| 16 | Pz | 23.2*** | P6 | 2.78 |
| 17 | P7 | 21.9*** | P4 | 2.04 |
| 18 | P9 | 19.9*** | P2 | 1.50 |
| 19 | P10 | 18.5*** | P3 | 1.46 |
| 20 | P4 | 17.8*** | P1 | 1.14 |
| 21 | P2 | 17.6*** | Pz | 0.77 |

**Table S3. Significance of the general visual response at the medial occipital ROI for each group of infants (i.e., type of stimuli).** Significance was estimated for each harmonic and for the summed response across harmonics using *Z*-scores calculated on the average across electrodes within the ROI, odor contexts and infants within groups. Significance was fixed at $Z > 1.64$ ($p < .05$, one-tailed, signal > noise). Asterisks indicate significance (*** $p < .001$).

| | | Nat Group | Simp Group |
|---|---|---|---|
| **Harmonic** | **Frequency (Hz)** | ***Z*-score** | ***Z*-score** |
| 1 | 6 | 88.3*** | 56.9*** |
| 2 | 12 | 117*** | 76.4*** |
| 3 | 18 | 39.5*** | 36.7*** |
| 4 | 24 | 26.4*** | 9.74*** |
| Sum | 6 to 24 | 113*** | 85.1*** |

**Table S4. Significance of the general visual response at the medial occipital ROI for each individual infant within each group.** Significance was estimated for the summed response of each infant, at each electrode within the ROI and for the mean response across electrodes, using *Z*-scores calculated on the average across odor contexts. *Z*-scores in bold indicate significance ($Z > 1.64$, $p < .05$, one-tailed, signal > noise).

| | **Nat Group** | | | | |
|---|---|---|---|---|---|
| **Infant** | **POz** | **O1** | **Oz** | **O2** | **mO** |
| **#01** | **9.32** | **16.93** | **14.02** | **12.44** | **22.90** |
| **#02** | **27.15** | **16.30** | **22.12** | **24.15** | **32.51** |
| **#03** | **34.61** | **54.31** | **44.92** | **44.89** | **53.23** |
| **#04** | **11.57** | **45.94** | **58.31** | **42.89** | **60.01** |
| **#05** | **8.77** | **29.49** | **32.00** | **25.70** | **35.95** |
| **#06** | **3.57** | **17.81** | **47.94** | **33.11** | **38.12** |
| **#07** | **27.10** | **27.23** | **26.64** | **25.23** | **32.75** |
| **#08** | **24.50** | **24.63** | **30.45** | **26.31** | **43.79** |
| **#09** | **12.72** | **17.78** | **40.11** | **47.40** | **50.65** |
| **#10** | **3.98** | **15.32** | **37.33** | **43.06** | **44.54** |
| **#11** | **10.01** | **29.29** | **39.02** | **25.74** | **34.17** |
| **#12** | **19.01** | **15.39** | **22.31** | **17.10** | **34.45** |
| **#13** | **9.67** | **17.06** | **14.29** | **7.42** | **19.26** |
| **#14** | -0.20 | **3.92** | **4.25** | **6.10** | **5.37** |
| **#15** | **1.66** | **4.78** | **14.02** | **10.22** | **9.88** |
| **#16** | **12.99** | **31.01** | **72.01** | **44.66** | **57.97** |
| **#17** | **11.75** | **18.36** | **23.15** | **13.35** | **32.78** |
| **#18** | **32.40** | **18.20** | **77.36** | **55.56** | **82.36** |
| **#19** | 0.71 | **10.95** | **9.39** | **5.38** | **14.09** |
| **#20** | **3.62** | **2.20** | **7.88** | **7.30** | **7.81** |
| **#21** | **12.10** | **5.94** | **9.54** | **10.19** | **19.87** |

| | **Simp Group** | | | | |
|---|---|---|---|---|---|
| **Infant** | **POz** | **O1** | **Oz** | **O2** | **mO** |
| **#01** | **22.55** | **12.11** | **27.59** | **27.17** | **43.45** |
| **#02** | 0.92 | **2.72** | 0.72 | **5.22** | **3.98** |
| **#03** | **5.40** | **4.44** | **9.70** | **9.82** | **11.35** |
| **#04** | **5.11** | **12.21** | **10.58** | **5.00** | **17.08** |
| **#05** | **6.07** | **8.73** | **17.67** | **7.68** | **13.42** |
| **#06** | -1.05 | 1.64 | **3.60** | **4.06** | **3.36** |
| **#07** | **5.81** | **6.53** | **3.63** | -2.24 | **5.88** |
| **#08** | **2.17** | **33.92** | **28.37** | **8.22** | **31.00** |
| **#09** | **2.69** | **9.24** | **11.70** | **15.61** | **11.98** |
| **#10** | **7.37** | **10.88** | **6.11** | **3.23** | **12.85** |
| **#11** | **5.85** | **4.01** | **6.86** | **16.71** | **10.64** |
| **#12** | 0.42 | **5.14** | **5.89** | **2.62** | **5.39** |
| **#13** | 0.37 | 1.53 | 0.70 | -0.50 | 1.25 |
| **#14** | **51.65** | **44.70** | **51.27** | **21.10** | **52.73** |
| **#15** | **2.82** | **21.01** | **35.35** | **28.19** | **30.70** |
| **#16** | **11.66** | **8.86** | **15.16** | **6.95** | **15.48** |
| **#17** | **11.63** | **29.93** | **17.64** | **7.50** | **27.75** |
| **#18** | **4.70** | **27.48** | **22.38** | **14.33** | **34.35** |
| **#19** | 0.31 | 0.61 | **3.13** | **4.55** | **3.74** |
| **#20** | **6.32** | **14.59** | **43.74** | **26.76** | **46.55** |
| **#21** | **11.09** | **17.61** | **26.09** | **31.07** | **29.54** |

**Table S5. Significance of the face-selective response at each ROI and for each group of infants (i.e., type of stimuli).** Significance was estimated for each harmonic and for the summed response across harmonics using *Z*-scores calculated on the average across electrodes within the ROIs, odor contexts and infants within the group. Significance was fixed at $Z > 1.64$ ($p < .05$, one-tailed, signal > noise). Asterisks indicate significance (* $p < .05$, ** $p < .01$, *** $p < .001$). (lOT: left occipito-temporal ROI, mO: medial occipital ROI, rOT: right occipito-temporal ROI, 3ROIs: mean of the three ROIs).

| | | Nat Group | | | | Simp Group | | | |
|---|---|---|---|---|---|---|---|---|---|
| **Harmonic** | **Frequency (Hz)** | **lOT** | **mO** | **rOT** | **3ROIs** | **lOT** | **mO** | **rOT** | **3ROIs** |
| 1 | 1 | 3.45*** | 0.27 | 1.81* | 2.09* | 8.52*** | 5.41*** | 7.99*** | 9.24*** |
| 2 | 2 | 1.50 | 0.80 | 0.06 | 1.20 | 6.14*** | 4.99*** | 4.68*** | 9.19*** |
| 3 | 3 | -0.34 | -0.78 | -1.37 | -0.89 | 5.01*** | 3.89*** | 6.77*** | 8.15*** |
| 4 | 4 | 2.59** | 0.85 | 1.05 | 2.19* | 2.93*** | 3.04** | 0.54 | 2.64** |
| 5 | 5 | -0.09 | -0.45 | -0.84 | -0.64 | 3.28*** | 6.69*** | 2.17* | 5.42*** |
| Sum | 1 to 5 | 3.64*** | 0.23 | 1.38 | 2.38** | 11.1*** | 9.51*** | 8.78*** | 14.7*** |

**Table S6. Significance of the face-selective response at each electrode of the ROIs for each individual infant within each group.** Significance was estimated for the summed response of each infant and at each electrode within the ROIs using *Z*-scores calculated on the average across odor contexts. *Z*-scores in bold indicate significance ($Z > 1.64$, $p < .05$, one-tailed, signal > noise). (lOT: left occipito-temporal ROI, mO: medial occipital ROI, rOT: right occipito-temporal ROI).

| | Nat Group | | | | | | | | | | | | | |
|---|---|---|---|---|---|---|---|---|---|---|---|---|---|---|
| | **lOT** | | | | | **mO** | | | | **rOT** | | | | |
| **Infant** | **P9** | **P7** | **PO7** | **PO5** | **O1** | **PO3** | **POz** | **PO4** | **Oz** | **P10** | **P8** | **PO8** | **PO6** | **O2** |
| **#01** | -1.36 | -1.13 | -1.32 | -0.37 | -0.10 | -0.52 | 0.01 | 0.21 | -0.97 | **1.90** | **1.79** | **1.98** | 0.61 | 0.30 |
| **#02** | -0.62 | -0.55 | -0.46 | -0.21 | -0.18 | 0.22 | -0.67 | 0.38 | 0.84 | 0.63 | 0.15 | 0.31 | 0.71 | 0.48 |
| **#03** | 1.25 | **2.19** | 0.95 | -0.64 | 0.63 | -0.90 | 0.63 | 0.10 | 0.43 | -0.44 | 0.03 | -0.20 | -0.11 | -1.03 |
| **#04** | 0.30 | 0.63 | -0.12 | 1.01 | 0.77 | **2.55** | **4.36** | -0.43 | -0.02 | 0.63 | 0.81 | -0.32 | -1.47 | -1.65 |
| **#05** | -0.20 | -0.36 | 0.23 | -1.20 | -0.27 | -0.66 | -1.03 | -1.17 | -0.47 | 0.55 | 0.64 | 0.41 | -0.98 | -0.87 |
| **#06** | -0.03 | 0.25 | **1.81** | -1.20 | 1.55 | -0.80 | -0.87 | 0.05 | 0.69 | -0.30 | 0.52 | -1.01 | 0.97 | -1.12 |
| **#07** | **3.10** | 0.69 | **3.69** | **2.26** | **3.65** | **2.02** | 1.03 | **2.20** | **2.90** | **4.08** | **4.44** | **2.59** | **2.84** | **1.70** |
| **#08** | 0.04 | 0.17 | **3.01** | -0.11 | 1.33 | -0.72 | -0.72 | 0.93 | 0.76 | 0.53 | 0.57 | -0.15 | -0.20 | 0.52 |
| **#09** | -2.37 | -0.35 | 1.27 | 1.11 | **2.25** | 0.37 | 0.31 | 1.58 | **2.04** | 0.66 | **1.94** | **4.19** | **3.71** | **2.74** |
| **#10** | 0.73 | 0.80 | **2.68** | **3.92** | 1.22 | **3.91** | 0.99 | -2.15 | -0.29 | **1.86** | 1.01 | -1.98 | -2.13 | -3.31 |
| **#11** | -0.56 | -0.23 | **1.98** | 0.40 | -2.02 | 0.02 | 1.21 | 0.13 | 0.15 | -0.03 | 0.80 | 0.18 | 1.07 | -1.43 |
| **#12** | **2.23** | 1.11 | 1.36 | 1.51 | 1.33 | -1.38 | -0.53 | 0.01 | **1.71** | 0.70 | 0.27 | -1.57 | -1.45 | -1.34 |
| **#13** | 1.05 | 0.42 | 0.45 | -0.28 | 0.41 | -1.05 | 0.00 | 0.90 | 0.25 | 0.77 | -1.10 | **2.50** | 0.35 | -0.81 |
| **#14** | **2.30** | -0.92 | -0.49 | -0.70 | -0.97 | -0.73 | 1.23 | -0.49 | -1.92 | 0.95 | -0.08 | 1.44 | -1.67 | -2.74 |
| **#15** | **4.03** | **2.18** | **6.36** | **3.35** | **3.44** | **3.09** | 0.24 | 0.85 | **1.85** | -0.75 | 0.08 | -0.23 | -0.18 | 1.47 |
| **#16** | **1.73** | -0.79 | 0.76 | 0.61 | 0.24 | 0.58 | -0.03 | 0.02 | 1.39 | -0.58 | -0.94 | 0.65 | 0.08 | 1.19 |
| **#17** | -1.01 | -2.05 | 1.20 | -2.59 | **3.02** | -0.40 | 0.18 | -0.76 | 0.88 | -3.56 | -1.65 | 0.96 | 1.07 | **2.24** |
| **#18** | 1.54 | 0.80 | 1.15 | 1.02 | 1.12 | 1.04 | 0.39 | 0.95 | -0.54 | 1.32 | 1.26 | 1.13 | 1.15 | 0.00 |
| **#19** | 0.60 | 0.08 | 0.37 | -0.51 | 1.08 | -2.02 | -1.35 | **1.84** | 0.44 | 1.08 | 0.03 | -0.01 | -0.68 | -0.83 |
| **#20** | -1.05 | -1.64 | 1.07 | -0.16 | 0.94 | -1.60 | -0.92 | -0.15 | -2.24 | 0.51 | 0.26 | 1.08 | 0.44 | -0.93 |
| **#21** | -0.22 | 1.64 | 0.54 | 0.30 | -0.69 | -0.39 | -1.92 | -0.04 | -3.31 | -1.42 | -0.79 | 0.69 | -1.16 | 0.58 |

**Simp Group**

| | lOT | | | | | mO | | | | rOT | | | | |
|---|---|---|---|---|---|---|---|---|---|---|---|---|---|---|
| **Infant** | **P9** | **P7** | **PO7** | **PO5** | **O1** | **PO3** | **POz** | **PO4** | **Oz** | **P10** | **P8** | **PO8** | **PO6** | **O2** |
| #01 | 1.62 | -0.14 | 1.25 | 1.41 | 1.52 | **2.15** | 0.49 | 0.48 | **1.98** | **4.86** | **6.71** | **2.25** | **1.72** | **2.40** |
| #02 | -1.21 | -0.36 | -0.59 | -0.70 | 0.73 | 0.47 | -0.09 | -0.24 | 0.96 | -0.30 | -0.68 | -0.42 | **2.20** | -0.57 |
| #03 | 1.06 | -0.48 | 0.04 | 0.13 | -0.92 | 0.02 | 0.36 | 1.36 | -0.52 | **2.83** | **3.83** | 1.21 | 1.38 | -0.19 |
| #04 | **2.06** | -0.18 | 1.28 | **2.23** | -1.01 | 1.33 | -1.11 | 0.93 | -1.68 | 0.31 | -2.69 | -1.34 | -0.48 | -0.81 |
| #05 | **2.36** | **2.02** | **3.52** | **3.18** | **3.94** | **4.93** | 0.83 | 0.10 | **1.85** | 0.97 | **2.04** | -0.09 | -0.24 | -1.63 |
| #06 | 1.57 | **3.13** | **1.87** | **2.86** | **2.50** | **2.98** | 0.71 | 1.59 | **3.03** | 1.35 | **3.46** | **2.73** | **2.28** | **1.99** |
| #07 | -1.19 | 0.57 | 0.53 | 0.26 | 0.16 | 1.22 | 0.12 | **3.18** | 0.01 | 0.96 | 0.50 | **2.89** | **2.92** | 1.36 |
| #08 | 0.98 | -0.69 | 1.09 | 0.75 | **1.71** | -1.23 | -0.12 | 0.05 | 1.25 | 0.80 | 0.87 | -1.31 | -1.59 | 1.32 |
| #09 | 1.54 | 0.99 | 0.14 | 0.43 | -0.08 | 1.26 | 1.12 | -0.21 | -0.14 | 0.17 | -0.08 | 0.66 | 1.03 | -0.27 |
| #10 | **3.98** | 1.06 | 0.38 | **1.74** | **1.92** | 1.05 | 0.76 | **1.92** | **1.89** | 0.05 | -0.45 | -0.24 | 1.29 | **2.12** |
| #11 | 1.33 | 0.26 | -1.34 | 1.02 | 0.48 | **3.21** | **2.22** | **3.87** | 0.11 | 1.11 | **3.76** | **3.11** | **3.54** | 0.20 |
| #12 | **4.93** | **2.45** | -0.99 | 0.74 | -0.72 | 0.96 | -0.76 | -0.47 | **2.07** | 1.46 | **2.27** | -0.28 | -0.30 | 1.14 |
| #13 | **3.22** | 0.07 | -0.55 | 0.00 | 0.80 | -0.07 | -1.19 | 0.34 | -0.70 | -1.58 | -2.00 | -3.10 | 0.23 | -0.96 |
| #14 | 0.75 | 0.04 | 1.23 | 1.32 | 0.39 | -0.06 | 1.49 | 0.58 | 0.42 | 1.57 | 1.13 | **2.41** | 1.56 | 1.40 |
| #15 | 1.52 | 0.54 | **5.65** | **4.78** | **12.85** | **3.61** | **4.11** | **10.85** | **9.73** | **3.46** | **3.80** | **11.62** | **12.49** | **10.90** |
| #16 | **4.76** | **5.25** | **4.87** | **4.57** | **2.06** | **4.45** | 1.25 | **1.71** | **2.49** | **8.16** | **2.07** | **5.58** | 1.44 | **3.59** |
| #17 | **4.63** | 1.15 | **3.26** | **5.19** | 1.22 | **3.31** | 1.45 | **2.27** | **1.69** | **5.35** | -0.30 | **1.94** | **2.27** | **2.30** |
| #18 | **6.91** | **6.66** | **4.13** | 0.63 | **6.25** | 0.02 | 1.04 | **2.26** | **2.42** | **2.62** | 0.38 | 1.09 | 0.30 | **2.56** |
| #19 | 0.88 | 1.53 | 1.10 | -0.11 | **3.41** | -0.09 | 1.09 | -1.75 | -0.63 | **2.14** | -1.47 | 1.51 | -1.05 | 0.38 |
| #20 | 0.59 | 0.42 | **2.18** | **1.97** | -0.06 | 1.33 | 0.71 | 0.30 | -0.95 | 0.13 | -0.93 | -0.89 | -0.89 | -1.52 |
| #21 | **5.26** | **4.69** | **6.05** | **6.96** | **3.27** | **3.92** | **2.14** | **1.82** | **4.03** | **18.66** | **6.88** | **11.15** | **2.35** | 0.90 |

**Table S7. ANOVAs and ANCOVAs conducted on the general and face-selective visual responses averaged across odor contexts.** For each response, individual amplitudes of the summed response across harmonics were submitted to a repeated-measures ANOVA using *Group* as a categorical between-subject factor. For the face-selective response, *ROI* was also used as a categorical within-subject factor. To account for the potential effect of the number of averaged segments per infant (*N Segments*), an additional ANCOVA was conducted using this variable as a continuous factor together with the categorical factors. Significant effects are indicated in red and values in brackets were obtained after application of the Greenhouse-Geisser correction for sphericity violation (as estimated by Mauchly's tests, unreported). SS: Sum of squares, Df: Degrees of freedom, MS: Mean square, $\eta_p^2$: partial eta squared.

**General visual response**

| ANOVA | SS | Df | MS | *F* | *p* | $\eta_p^2$ |
|---|---|---|---|---|---|---|
| *Group* | 51.2 | 1 | 51.2 | 7.47 | .009 | .16 |
| Error | 274.3 | 40 | 6.86 | | | |

| ANCOVA | SS | Df | MS | *F* | *p* | $\eta_p^2$ |
|---|---|---|---|---|---|---|
| *N Segments* | 3.69 | 1 | 3.69 | 0.53 | .47 | .01 |
| *Group* | 54.9 | 1 | 54.9 | 7.91 | .008 | .17 |
| Error | 270.6 | 39 | 6.94 | | | |

**Face-selective response**

| ANOVA | SS | Df | MS | *F* | *p* | $\eta_p^2$ |
|---|---|---|---|---|---|---|
| *Group* | 85.3 | 1 | 85.3 | 14.9 | .0004 | .27 |
| Error | 228.7 | 40 | 5.72 | | | |
| *ROI* | 8.27 | 2 (1.5) | 4.14 | 3.91 | .024 (.035) | .09 |
| *Group* × *ROI* | 0.19 | 2 (1.5) | 0.09 | 0.09 | .91 (.86) | .00 |
| Error | 84.6 | 80 (61.5) | 1.06 | | | |

| ANCOVA | SS | Df | MS | *F* | *p* | $\eta_p^2$ |
|---|---|---|---|---|---|---|
| *N Segments* | 6.73 | 1 | 6.73 | 1.18 | .28 | .03 |
| *Group* | 91.9 | 1 | 91.9 | 16.2 | .0003 | .29 |
| Error | 221.9 | 39 | 5.69 | | | |
| *ROI* | 0.36 | 2 (1.5) | 0.18 | 0.17 | .85 (.79) | .00 |
| *Group* × *ROI* | 0.14 | 2 (1.5) | 0.06 | 0.06 | .94 (.89) | .00 |
| Error | 84.4 | 78 (60) | 1.08 | | | |

**Table S8. ANOVAs and ANCOVAs conducted on the general and face-selective visual responses dissociated across odor contexts.** For each response, individual normalized amplitudes of the summed response across harmonics were submitted to a repeated-measures ANOVA using *Group* as a categorical between-subject factor and *Odor* as a categorical within-subject factor. For the face-selective response, *ROI* was also used as a categorical within-subject factor. Only effects involving the *Odor* factor are reported. To account for the potential effect of the number of averaged segments per infant (*N Segments*), an additional ANCOVA was conducted using this variable as a continuous factor together with the categorical factors. Significant effects are indicated in red. SS: Sum of squares, Df: Degrees of freedom, MS: Mean square, $\eta_p^2$: partial eta squared.

**General visual response**

| ANOVA | SS | Df | MS | *F* | *p* | $\eta_p^2$ |
|---|---|---|---|---|---|---|
| *Odor* | 0.00 | 1 | 0.00 | 0.00 | .94 | .00 |
| *Odor* × *Group* | 0.66 | 1 | 0.66 | 1.78 | .19 | .04 |
| Error | 14.9 | 40 | 0.37 | | | |

| ANCOVA | SS | Df | MS | *F* | *p* | $\eta_p^2$ |
|---|---|---|---|---|---|---|
| *N Segments* | 0.17 | 1 | 0.17 | 0.08 | .77 | .00 |
| Error | 79.8 | 39 | 2.05 | | | |
| *Odor* | 0.19 | 1 | 0.19 | 0.53 | .47 | .01 |
| *Odor* × *Group* | 0.82 | 1 | 0.82 | 2.19 | .15 | .05 |
| Error | 14.6 | 39 | 0.38 | | | |

**Face-selective response**

| ANOVA | SS | Df | MS | *F* | *p* | $\eta_p^2$ |
|---|---|---|---|---|---|---|
| *Odor* | 5.04 | 1 | 5.04 | 2.68 | .11 | .06 |
| *Odor* × *Group* | 9.78 | 1 | 9.78 | 5.19 | .028 | .11 |
| Error | 75.4 | 40 | 1.88 | | | |
| *Odor* × *ROI* | 7.42 | 2 | 3.71 | 3.14 | .049 | .07 |
| *Odor* × *Group* × *ROI* | 1.31 | 2 | 0.66 | 0.55 | .58 | .01 |
| Error | 94.6 | 80 | 1.18 | | | |

| ANCOVA | SS | Df | MS | *F* | *p* | $\eta_p^2$ |
|---|---|---|---|---|---|---|
| *N Segments* | 14.8 | 1 | 14.8 | 2.56 | .12 | .06 |
| Error | 225.2 | 39 | 5.78 | | | |
| *Odor* | 1.82 | 1 | 1.82 | 0.95 | .34 | .02 |
| *Odor* × *Group* | 10.3 | 1 | 10.3 | 5.36 | .026 | .12 |
| Error | 74.9 | 39 | 1.92 | | | |
| *Odor* × *ROI* | 7.45 | 2 | 3.73 | 3.26 | .044 | .08 |
| *Odor* × *Group* × *ROI* | 1.61 | 2 | 0.80 | 0.70 | .49 | .02 |
| Error | 89.1 | 78 | 1.14 | | | |
